# Temporal and spatial distribution of microbial community in urban river in Dhaka

**DOI:** 10.1101/2024.11.26.625349

**Authors:** Md. Masud Alom, Aksa Hossain Nizhum, Ayman Bin Abdul Mannan, Munawar Sultana, Sheikh Mokhlesur Rahman

**Author notes:** Corresponding Author: 1. Sheikh Mokhlesur Rahman, Associate Professor, Department of Civil Engineering, Bangladesh University of Engineering and Technology. Co-corresponding Author: Munawar Sultana, Professor, Department of Microbiology, University of Dhaka,.

## Abstract

Buriganga River in central Bangladesh represents a complex ecosystem with extreme pollution and altered water quality. It is well known that microorganisms play a key role in regulating the function and health of aquatic ecosystems. Microbial activities also have a direct impact on biogeochemical cycling and degradation of pollutants. Despite numerous studies on the physicochemical parameters of the water quality of this central river, a comprehensive understanding of microbial diversity and the impact of water’s physicochemical properties on bacterial community structure is not well documented. Therefore, the present investigation was done to elucidate the spatial and temporal shifts of microbial communities in the Buriganga River water in correlation to different physicochemical factors as well as heavy metal content. A total of 13 distinct sampling sites (from Amin Bazar Bridge to Postogola Cantonment B-C Friendship Bridge) encompassing a length of approximately 19 km along the river’s centre were selected to gather the sample seasonally during pre-monsoon, monsoon, post-monsoon, and winter. 16S rRNA gene-based metagenomics analysis revealed significant variation of bacterial diversity in winter rather than the rest of the three seasons with minimal variation suggesting stable geochemical cycles. Also, microbial diversity was higher at sampling points downstream compared to upstream along the river with overall dominance of *Proteobacteria* and *Firmicutes*. Physicochemical water quality parameters analysis identified Zn, TP (Total phosphate), BOD, and COD as key factors influencing bacterial community diversity and composition, although they were not the sole contributors. At the heavily polluted Lohar Bridge (10CS) site, significant nitrogen cycle disruption was evident. The high abundance of metal and antibiotic-resistant bacteria genera such as *Pseudomonas*, *Ralstonia*, and *Acinetobacter* at specific sampling positions indicated a relation between antibiotic and heavy metal pollution from adjacent hospitals and industrial effluent release points. The prevalence of metabolically unique bacteria such as *Geobacter* and *Dehalococcoides*, that are well known for utilizing specific heavy metals like iron and uranium as well as chlorinated compounds, might indicate the selective pressure of metal pollution from nearby working factories/industries. The investigation also identified potential pathogens like *Arcobacter, Acinetobacter,* and *Pseudomonas,* posing potential risks to both the environment and human health. Therefore, the Buriganga River belt serves as a potential hub for metal-antibiotic tolerant/resistant species with a direct influence on indigenous river-flowing communities correlating unique nutrient/geochemical cycling. The overall microbial community reflects unrestricted pollution in Buriganga central river aggravating water quality for regular use, indicating urgent intervention.

**GRAPHICAL ABSTRACT:** 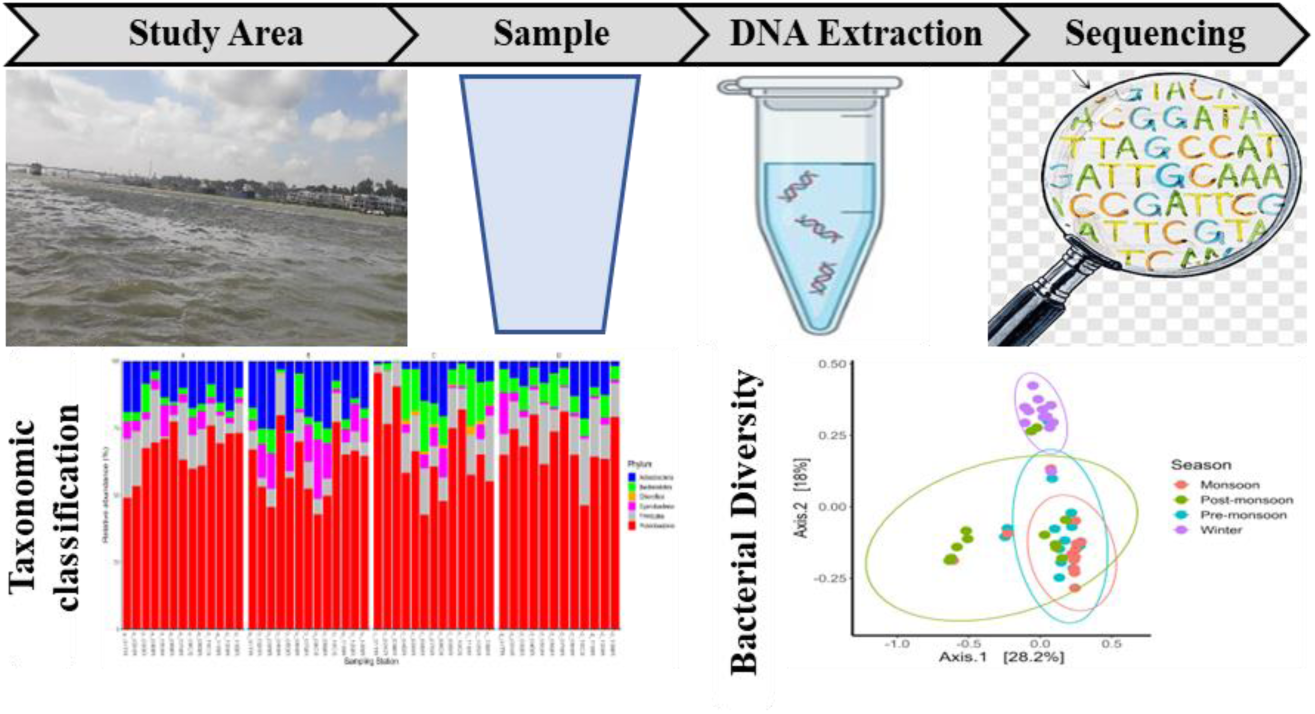

**HIGHLIGHT:** *Significant bacterial diversity was observed during winter compared to the other three seasons (Premonsoon, monsoon, and post monsoon)*.

*Zn, TP, BOD, and COD were found to be key factors influencing the diversity and composition of the bacterial community*.

*Disruption of the nitrogen cycle was evident at the heavily polluted monitoring point*.

*Potential pathogens were identified that may pose risks to both the environment and human health*.

## 1. Introduction

Urban rivers are a part of the aquatic environment which plays a vital role to supply water for human consumption and industrial activities especially those along the river (Breton-Deval et al., 2019; Zhang et al., 2021). These rivers also carry stormwater run-off and urban sewage (Wang et al., 2018) comes from the various locations of the city. Rapid industrial growth and various human activities lead to increase in the concentration of pollutants like phosphorus, organic matter and heavy metals in rivers. The increasing pollution level in the river degrades the aquatic ecosystems (Wu et al., 2018; Huang F. et al., 2019; Qin et al., 2020, 2021; Sheng et al., 2013; Liang et al., 2018). It is indicated that the discharge of wastewater from diverse industries plays a key factor in breaking the ecological balance of urban rivers (Raat, 2001; Rajaram and Das, 2008). Microorganisms that are living in these ecosystems are crucial for maintaining ecological balance and transforming pollutants through participation in nutrient and organic matter cycle processes (Bai et al., 2014; Ruiz-González et al., 2015). Fluctuations in physicochemical water parameters due to pollution can impact microbial diversity and community structure (Kostanjsek et al., 2005; Lundgaard et al., 2017; Tiquia, 2010).

Microorganisms have significant roles in purifying the water as well as the formation of black-odor rivers (Liang et al., 2021; Sun et al., 2018). It is proved that microbial activities have a direct impact on biogeochemical cycling and degradation of pollutants. It also has a great effect on regulating the ecological function, and sustaining the health of aquatic ecosystems (Chen et al., 2019; Li et al., 2021a). Furthermore, microorganisms have an influence on changing the river water condition e.g. high/low amounts of organic matter, heavy metals, or nutrients etc. (Chen et al., 2017; Wang et al., 2021). It is pointed out in different research that, for the disturbance of physicochemical parameters, microbial community structure and metabolism have forthright reflection (Gao et al., 2021; Zhang et al., 2020a; Zhang et al., 2019a, 2019b; Zhang et al., 2020b). The bacterial community structure can be reshaped by the alteration of diverse parameters such as pH, temperature, phosphorus, dissolved oxygen content etc. The change of those parameters can be done by containing organic and inorganic pollutants which may be added in river water from diverse sources like sewage from the residential and industrial areas. (Garcia-Armisen et al., 2014; Lindström et al., 2005; Mark Ibekwe et al., 2012; Wang et al., 2018). Therefore, to maintain a sustainable healthy ecosystem in an urban river, it is urgent to monitor the correlation between water quality parameters and microbial communities from time to time. Some research has already been done where researchers reported the seasonal effect on the microbial community compositions in rivers. Zhang et al., (2019a, 2019b) and Zhu et al., (2019) mentioned in their studies that the seasonal effects on the microbial community structures are greater than the locations. On the other hand, Li et al., (2019) got different results in their observation. So, temporal and spatial investigations are needed for a polluted river to understand the bacterial community structure properly. These might help to get an idea about the degree of contamination and the function of the river ecosystem with seasonal and environmental change. Besides, it may help to detect the potential pathogens and novel organisms which are responsible for bioremediation.

The Buriganga River which flows through the southern part of Dhaka city, the capital of Bangladesh (Saifullah et al., 2013; Rahman and Al Bakri, 2010). Unfortunately, the current state of this river is dire, marking it as the most polluted river in Bangladesh and facing a precarious existence. This dire condition results from numerous factors, including waste from mills and factories, household refuse, medical waste, animal carcasses, plastics, oils, and various other pollutants (Tamim et al., 2016; Islam and Azam, 2015). Previous studies have primarily focused on the impact of urban activities on the river’s surface water and sediments, emphasizing physicochemical indices such as heavy metals and nutrients (Islam et al., 2014; Bhuiyan et al., 2015). However, there has been a noticeable gap in research concerning the river’s microbial community, in contrast to the extensive assessments of its water (Fatema et al., 2018) and sediment (Tamim et al., 2016) properties.

To our best knowledge, understanding of microbial community composition and diversity in the Buriganga River water is still missing. Thus, for a better understanding of the bacterial community in water at different sites and seasons, we performed 16S rRNA gene-based high throughput sequencing on the microbial communities in this study. Analyzing microbial communities through the utilization of 16S rRNA genes represents a direct and economical approach to characterize the taxonomic composition of a microbial community. However, its taxonomic resolution is constrained by the preservation of the target gene and the length of the amplicon product. The dependency on the 16S rRNA gene also restricts our capacity to profile non-bacterial constituents in the microbial community, including Archaea and Eukarya. Additionally, it has challenges to determine the functional capabilities of the microorganisms (Douglas et al., 2020; Wemheuer et al., 2020; Peterson et al., 2021). The main objectives of this study are to explore the taxonomic diversity of existing bacterial communities at selected sites of the Buriganga River belt and in varying seasons and to correlate the environmental factors for shifting of the bacterial community.

## 2. Materials and Methods

### 2.1 Study Area

The Buriganga River, flowing along the southern and western peripheries of Dhaka city, is a tidal watercourse sourced from the Dhaleshwari River through the Karanatali and Turag tributaries. The Buriganga River holds significant economic importance for Dhaka city, serving as a vital waterway for river transportation via launches and country boats. A total 13 monitoring stations were judiciously chosen along the Buriganga River and its tributaries to assess the diversity. Among these, 8 stations were along the Buriganga River which were selected based on zones where the river receives significant amounts of residential and industrial wastewater, two points (08CS and 10CS) were taken inside the canals connected to the river, chosen primarily to assess the impact of canal water—which passes through numerous residential areas—on the river’s water quality. Additionally, two sites were established upstream where two other rivers (Turag, 01TR and Karnatali, 02KR) converge with the Buriganga. This was done to understand variations in water quality and microbial activity in these areas. A nother station 03BR was selected at the confluence of three rivers (Buriganga, Turag, and Karnatali) (Fig. 1). Details about the water quality monitoring stations can be found in Table S1.

**Fig. 1.**
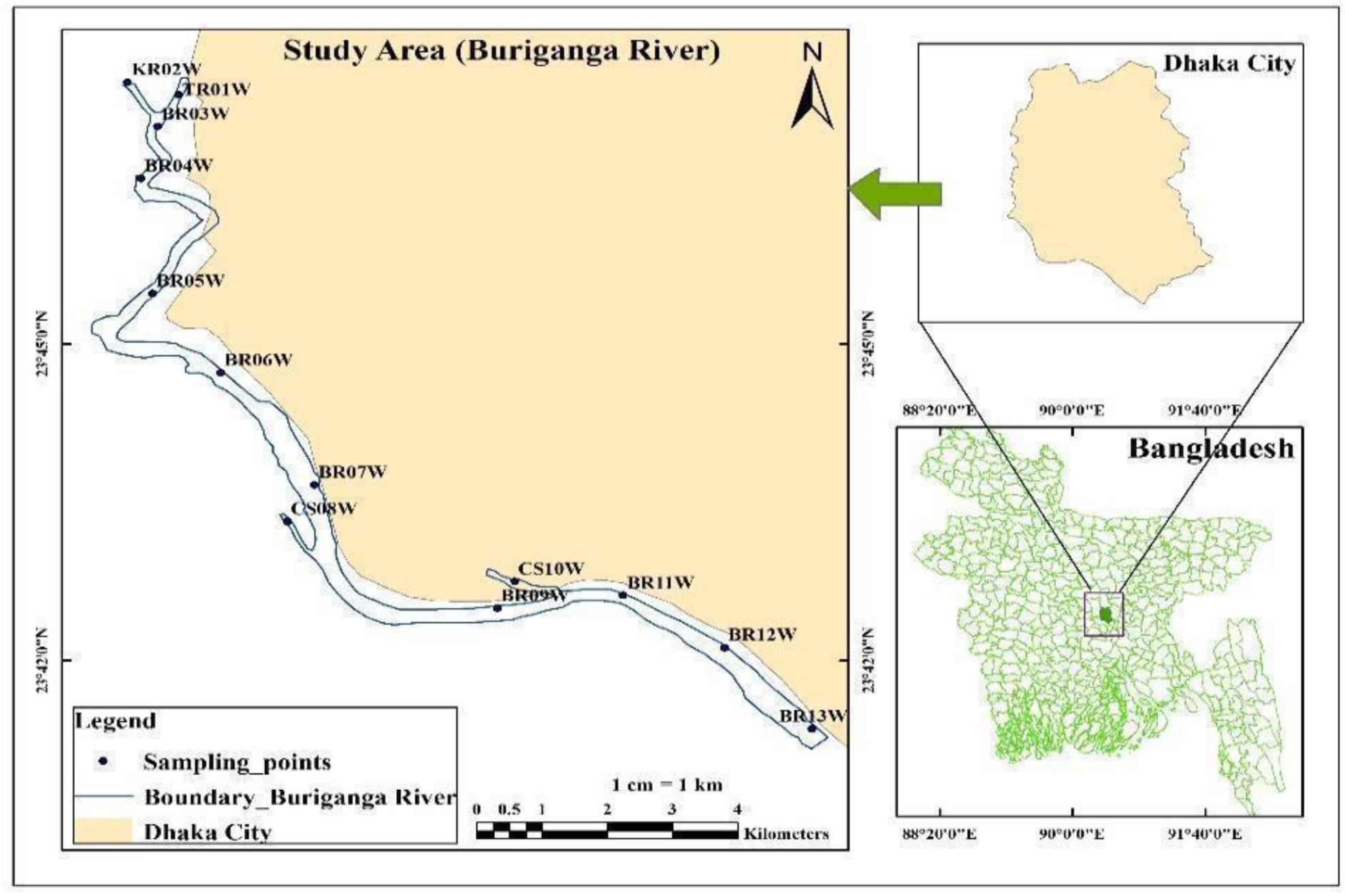
Detailed map of the Study area (generated by using Google Earth Pro and GIS software).

### 2.2 Sample Collection and Physicochemical Characteristics Analysis

Water samples were obtained from 13 sampling stations along the Buriganga River, spanning four distinct seasons: Pre-monsoon (May 2022), Monsoon (August 2022), Post-monsoon (November 2022), and Winter (January 2023). Positioned 10 meters from the shore, samples were manually collected at a depth of 15 cm using plastic bottles of varying sizes (2 L and 500 ml) (Dong et al., 2023). Within one hour of collection, the samples were promptly transported to the laboratory and preserved at 4°C to facilitate subsequent analysis within two days. To mitigate the risk of unintended contamination, bottles were meticulously rinsed three times with sampling water before collecting samples at each station (Chen et al., 2007). A total of 18 water quality parameters were examined for each sample, including pH, Turbidity, Electric Conductivity (EC), Total Dissolved Solids (TDS), Temperature, Dissolved Oxygen (DO), Biochemical Oxygen Demand (BOD5), Chemical Oxygen Demand (COD), Nitrogen Dioxide (NO2), Ammonium Nitrate (NH3-N), Total Phosphorus (TP), Chloride (Cl-), Lead (Pb), Cadmium (Cd), Chromium (Cr), Copper (Cu), Nickel (Ni), and Zinc (Zn). Among these parameters, pH, DO, Temperature, and Turbidity were promptly determined at the sampling site, while the rest were analyzed at the Laboratory. Detailed information regarding the analytical methods employed for each parameter can be found in Table S2.

### 2.3 Extraction of DNA from Water Samples

The genomic DNA from each water sample was extracted by using the DNeasy® PowerWater® Kit (Qiagen, USA). For this, 250 ml of water sample from each sampling site was filtered with the help of a filter funnel attached with a vacuum source. A filter paper with 0.20 µm was used in this process. After filtering, the filter paper with the microbial load was removed in an aseptic condition and it was inserted into a 5 ml Power Water Bead Pro tube which was provided with the Kit box and followed the manufacturer’s protocol (DNeasy PowerWater Kit Handbook, 2022) for further steps to complete the extraction process. The quantity and purity were determined with Nanodrop (Thermosphere, USA) by measuring 260/280 absorbance ratios. The extracted DNA was stored at -40 ^0^C until use (Wang et al., 2018; Zhang et al., 2021).

### 2.4 Illumina Miseq Sequencing of 16S rRNA Gene Amplicons

Illumina Miseq metagenomic sequencing of 16S rRNA gene was employed to observe the composition of the bacterial community. The primers 341F and 806R (barcode- CCTAYGGGRBGCASCAG and barcode-GGACTACNNGGGTATCTAAT) were used to amplify the V3-V4 hypervariable regions of the bacterial 16S ribosomal RNA gene to each sample (Ouyang et al., 2020).

### 2.5 Taxonomy Identification

The Miseq sequencing data were analyzed using the QIIME 2 (Quantitative Insights Into Microbial Ecology 2, version 2022.8) software package (Caporaso et al., 2010b; Wang et al., 2018). Raw sequences were processed by eliminating quality scores below 20 for further analysis. The data was de-noised by using DADA2 tool (Peterson et al., 2021; Callahan et al., 2016). Finally, high-quality sequences were clustered into Operational Taxonomic Units (OTUs) based on 97% sequence similarity with GreenGenes (v13_8_99) (Sierra et al., 2020; McDonald et al.,2012). Utilizing Illumina MiSeq Next Generation Sequencing (NGS), a total of 4,956,751 reads were generated from the 51 samples, averaging 97,191 reads per sample (Table S3). These reads were categorized into 2098 Operational Taxonomic Units (OTUs). The rarefaction curves, depicting species against read numbers (Figure S1) indicated ample sequencing depth.

### 2.6 Statistical Analysis

Statistical analyses were carried out in R (version 4.3.1) and RStudio-2023.06.1-524 interface. Rarefaction curves were generated to assess the completeness of sampling. Analyses of bacterial community composition were carried out using the ’vegan’ package in R. Hierarchical cluster analysis (CA) was utilized to assess the similarities and differences among the sampling points in the dataset. Ward’s method was applied to evaluate the similarity among the monitoring sites (Zhao et al., 2012). For comparing the richness and diversity of bacterial communities in each sample, alpha diversity indexes such as the Shannon index and Simpson index were computed. Bacterial α-diversity was calculated post-normalization of sequencing depth, utilizing the sample with the smallest sequencing effort as the basis. To assess β-diversity among various samples, a Principal Coordinates Analysis (PCoA) of Bray–Curtis’s distance, along with a PERMANOVA test (Permutational Multivariate Analysis of Variance) (Xie et al., 2021), was performed. Seasonal variations were examined using the Wilcox.test, where a p-value significance threshold of 0.05 was employed (Wang et al., 2018). For finding the common phylum and genera among the four seasons Venn diagram was plotted by using VEENY 2.1 (https://bioinfogp.cnb.csic.es/tools/venny/) tool.

## 3. Result

### 3.1 Seasonal Bacterial Profile in the River Water

#### Abundances

Bacterial community composition and structure in the water of the Buriganga River exhibited seasonal and spatial variations at both phylum and genus levels. Figure 2 illustrates the dominant phyla (Fig. 2A) and genera (Fig. 2B) present in the water samples across different seasons. On average, across the four seasons, the phylum, *Proteobacteria* was the most dominant, comprising 65.72% of the community, followed by *Actinobacteria* (11.53%), *Firmicutes* (11.33%), *Bacteroides* (6.32%), and *Cyanobacteria* (4.86%). At the genus level, *Acinetobacter* was the most prevalent, accounting for 20.82% of the community, followed by *C39* (*Rhodocyclaceae*_Family) (12.46%), *Arcobacter* (11.72%), *Novosphingobium* (9.42%), *Dechloromonas* (8.09%), and *Hydrogenophaga* (6.74%), among others (Table S4 and S5).

**Fig. 2.**
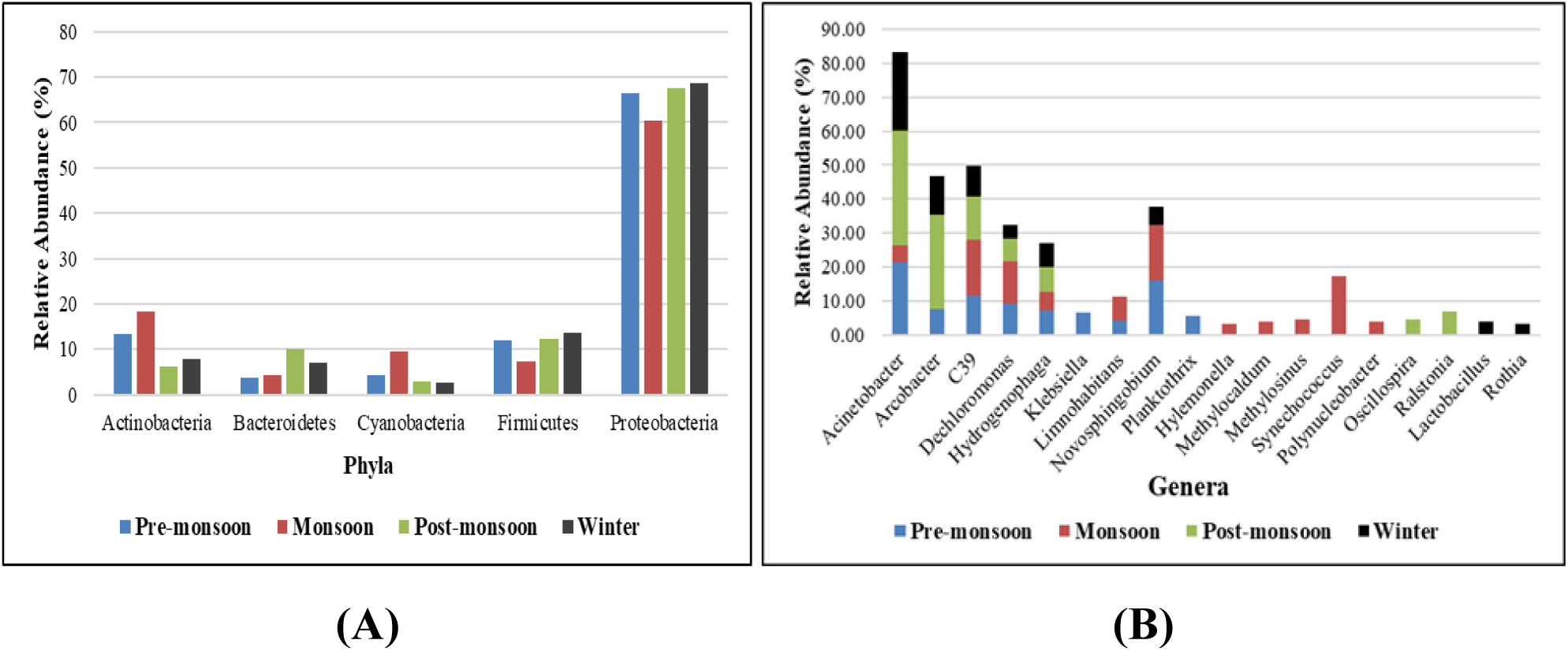
Seasonal variation of different Phyla **(A)** and Genera (RA > 3%) **(B)** among the four distinct seasons with respect to their relative abundance (%)

During the pre-monsoon season, *Proteobacteria* showed a high prevalence, with an average relative abundance of 66.41% across different sites. Previous studies have indicated that *Proteobacteria* are capable of surviving in stressful environments (Bååth & Kritzberg, 2024; Spain et al., 2009). *Actinobacteria* also exhibited the second-highest abundance (13.5%) in this season due to their resilience in warmer conditions (20°C to 40°C) (Sreevidya et al., 2016). Water temperature in the Buriganga River is observed to range from 29.2°C to 30.4°C, consistent across various sites at this time (Alom et al., 2024_Preprint). At the genus level, *Acinetobacter* (21.32%) and *Novosphingobium* (15.98%) were prominent, underscoring their adaptability to higher temperatures.

During the monsoon season, *Proteobacteria* continued to dominate, with an average relative abundance of 60.4%, likely due to the influx of rainwater, bringing more organic material and nutrients into the water (Wen et al., 2023). *Actinobacteria* and *Cyanobacteria*, known for thriving in nutrient-rich and light-abundant environments, also showed higher relative abundances 18.39 % and 9.58% respectively. At the genus level, *Synechococcus* (17.42%), *Novosphingobium* (16.42%) and *C39* (Rhodocyclaceae Family) (16.34%) were particularly notable during this season.

In the post-monsoon, *Proteobacteria* remained highly abundant (67.49%), similar to other seasons. *Firmicutes* showed a notable increase, rising from 7.25% in the monsoon to 12.39% in the post-monsoon, indicating their adaptation to the organic matter decomposition typical of this season.

*Bacteroidetes* also demonstrated a higher average abundance (10.05%) across the monitoring points. At the genus level, there was a significant increase in the average relative abundance of *Acinetobacter*, jumping from 4.97% in the monsoon to 34.03% in the post-monsoon. *Arcobacter* exhibited a similar trend, reaching 27.58% during this season.

*Proteobacteria* exhibited the highest abundance (68.57%) in winter compared to the other seasons, while *Firmicutes* continued to thrive, particularly in stable environments with sufficient organic matter. The presence of *Firmicutes* (13.6%) during this time underscores their resilience and ability to persist in colder conditions. *Cyanobacteria* (2.69%) had a lower abundance likely due to reduced light availability and cooler temperatures. It is recorded the range of winter temperatures in this study from 21.5°C to 23.4°C (Alom et al., 2024_Preprint). At the genus level, *Acinetobacter* (22.98%) predominated followed by *Arcobacter* (11.66%), *C39* (9.11%), *Hydrogenophaga* (6.87%) and so on.

#### Unique and shared microbial community among the four seasons

The Venn diagram (Fig. 3) illustrates the shared phyla and genera across the four seasons. Out of the total six phyla, five were found to be common throughout all four seasons (*Proteobacteria, Actinobacteria, Firmicutes, Bacteroidetes,* and *Cyanobacteria*) (Fig. 3A). Conversely, among the total 33 genera, only four were identified as common across the seasons (*Acinetobacter, Cupriavidus necator C39, Dechloromonas,* and *Hydrogenophaga*) (Fig. 3B). As depicted in Fig. 3B, during the winter season, 14 distinct genera were observed at selected sites, contrasting sharply with the bacteria found during other seasons (Pre-monsoon, Monsoon, and Post-monsoon). This variation could potentially be attributed to changes in water quality. During winter, the water quality of the Buriganga River tends to deteriorate compared to other seasons.

**Fig. 3.**
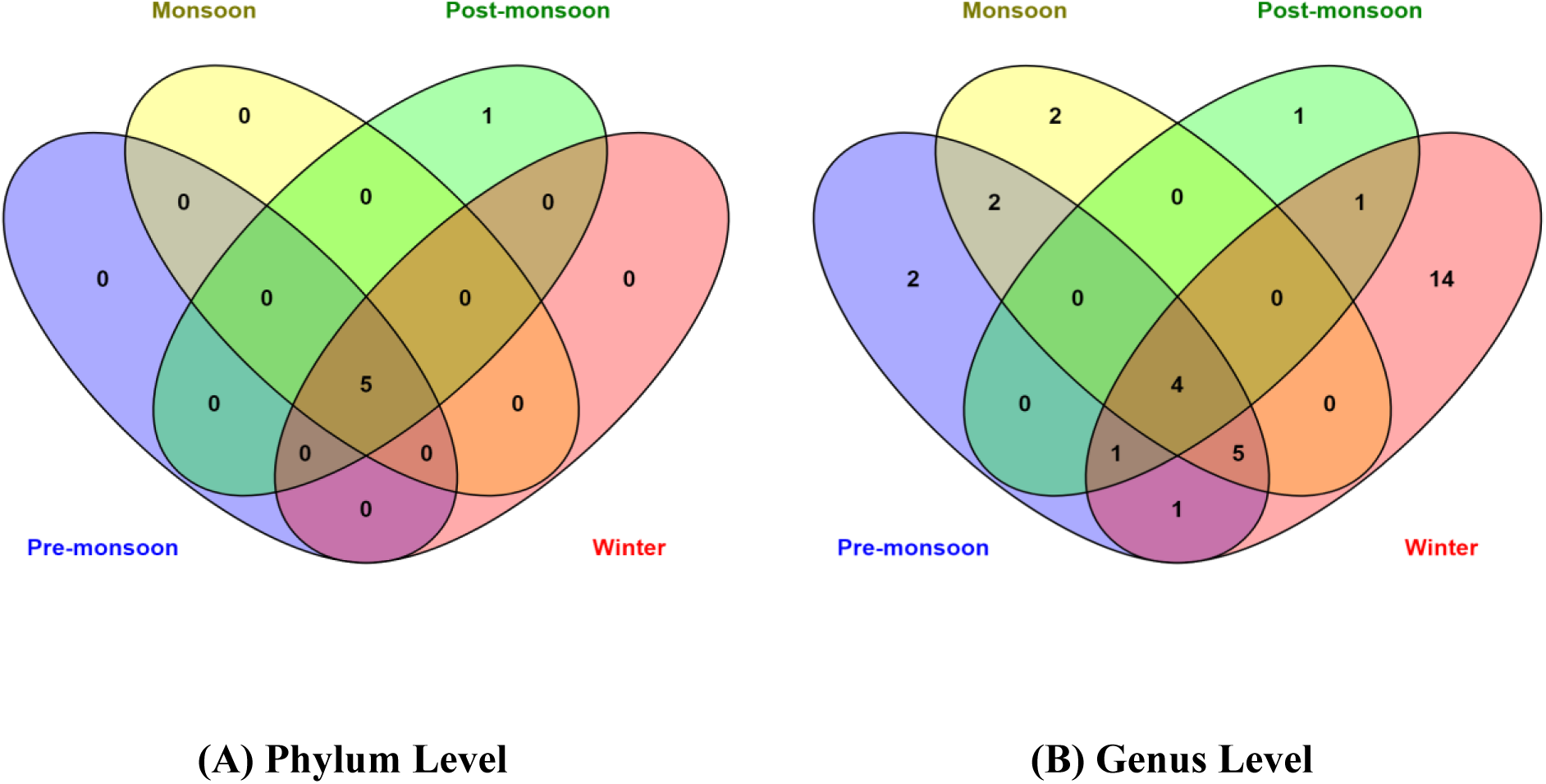
Venn diagram identifying the unique and shared microbial community among the four seasons at **(A)** phylum and **(B)** genus level (using tool VEENY 2.1 https://bioinfogp.cnb.csic.es/tools/venny/)

### 3.2 Spatial Changes of Bacterial Community

#### 3.2.1 At the phylum level

The analysis revealed six types of Phyla that were dominating in the water of the Buriganga River across the chosen 13 monitoring sites. *Proteobacteria* exhibited maximum abundance at every monitoring station in all four seasons, with the highest proportion (95.64%) observed at site 01TR during the post-monsoon period, and the lowest (31.42%) at site 06BR in the same season (Fig. 4). Prior studies have also pointed to *Proteobacteria* as the predominant microorganism in urban river ecosystems (Liu et al., 2015; Murphy et al., 2021; Shi et al., 2020). It can be mentioned that *Proteobacteria* is one of the most diverse and abundant groups of microbes on Earth. *Firmicutes* displayed maximum (24.54%) and minimum (0.77%) abundance at site 10CS (during winter) and 05BR (in the pre-monsoon), respectively. Research carried out on urban rivers in Mexico detected the presence of *Firmicutes* in regions influenced by human activity (Garcia-Mazcorro et al., 2016). Additionally, fluctuations in microbial community abundance can be influenced by environmental factors (Su et al., 2018). According to physicochemical parameters analysis, this station (10CS) is also identified as the most polluted site among the selected sites (Alom et al., 2024_Preprint). *Actinobacteria* (26.01%) and *Cyanobacteria* (21.98%) were the most abundant bacteria during the monsoon period at monitoring stations 05BR and 08CS, respectively, with the lowest abundances observed at stations 02KR and 06BR during the post-monsoon and monsoon periods. *Bacteroidetes* exhibited the highest (19.08%) and lowest (0.54%) abundances at site 11BR and 01TR, respectively, during the post-monsoon season (Fig. 4).

**Fig. 4.**
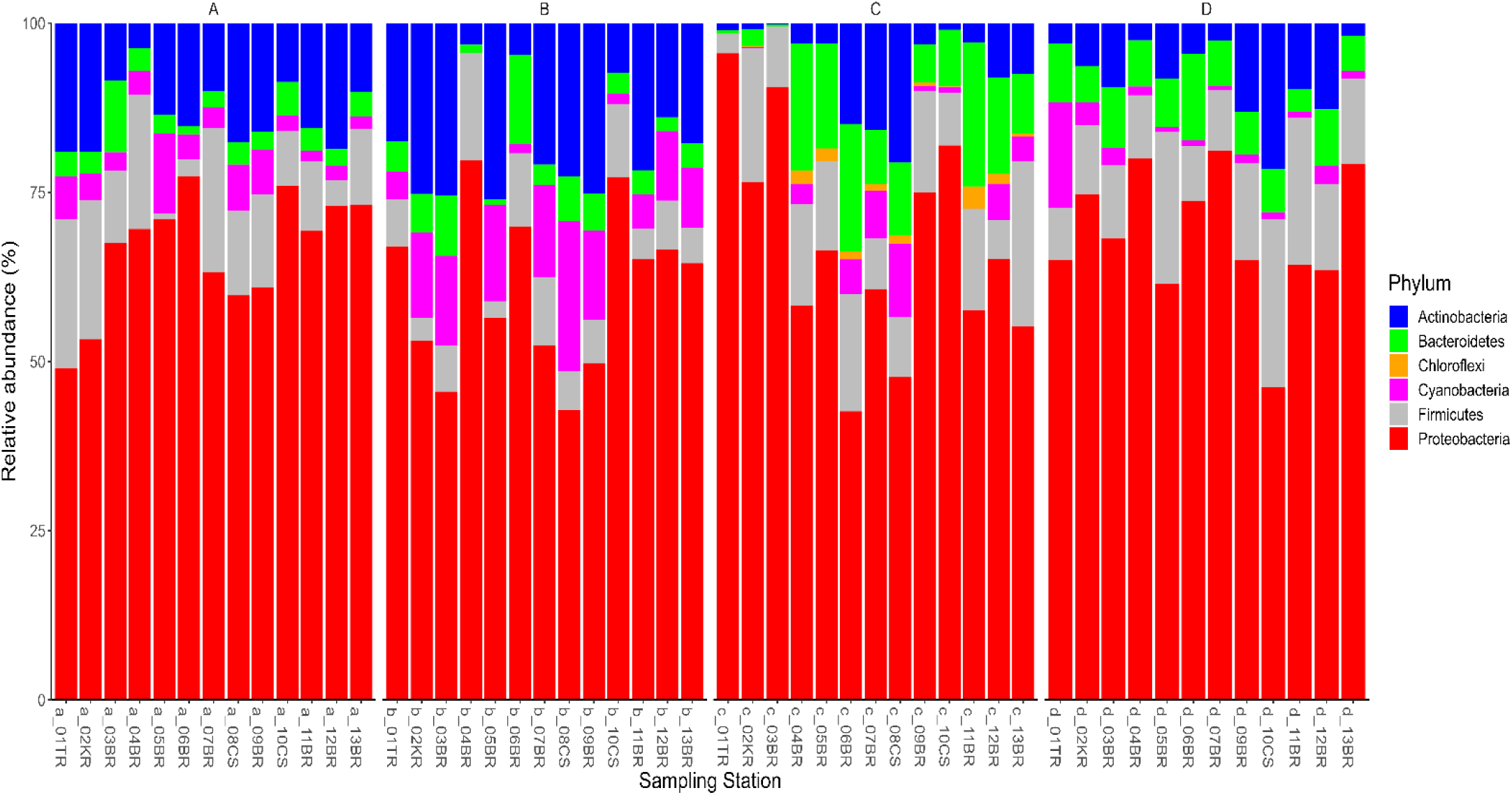
Taxonomic classification of the bacterial community (Phyla) at each site (consider relative abundance > 0.5%); A, B, C, and D indicate the pre-monsoon, monsoon, post-monsoon, and winter seasons accordingly.

#### 3.2.2 At the genus level

A total of 33 Genera were identified in this study, with approximately 6 being predominant. Changes in the abundance of various genera may occur due to variations in pollution intensity and fluctuations in water quality parameters. Additionally, the flow patterns of river water could be another factor affecting the abundance of microorganisms in different sections of the river. Stations 01TR and 02KR, situated inside the Turag and Karnatali rivers respectively, which are connected to the Buriganga River, exhibited almost similar bacterial compositions. Analysis revealed that (Fig. 5), at stations 01TR, 02KR, and junction point 03BR (where Buriganga, Turag, and Karnatali meet), with *Pseudomonas/Ralstonia* showing maximum abundance (around 95%) during the post-monsoon season. Other less dominant bacteria at these stations during the remaining seasons included *Acinetobacter, C39 (Rhodocyclaceae_*Family), *Synechococcus,* and so on. Stations 08CS and 10CS, located in the two different canals connected to the Buriganga River, exhibited varied bacterial abundances. *Arcobacter* dominated at station 10CS during the post-monsoon period (55.43%), while *Acinetobacter, Dechloromonas, Pseudomonas, C39 (Rhodocyclaceae_*Family), and *Hydrogenophaga* were also present with low abundance in different periods of the year. Conversely, *Acinetobacter* (33.43%) was the most abundant at site 08CS during the pre-monsoon season, accompanied by *C39 (Rhodocyclaceae_*Family) and *Synechococcus,* among others, in different seasons. Among the 13 stations, 8 were situated along the Buriganga River, where *Acinetobacter* was the most prevalent bacteria at sites 13BR (75.23%), 04BR (13.90%), 05BR (19.79%), and 06BR (17.72%) during winter. Additionally, *Acinetobacter* dominated at 09BR (55.34%) during the post-monsoon period. Less abundant bacteria such as *Lactobacillus, C39 (Rhodocyclaceae_*Family*), Novosphingobium, Synechococcus,* and *Geobacter* were also detected across those stations in different seasons.

**Fig. 5.**
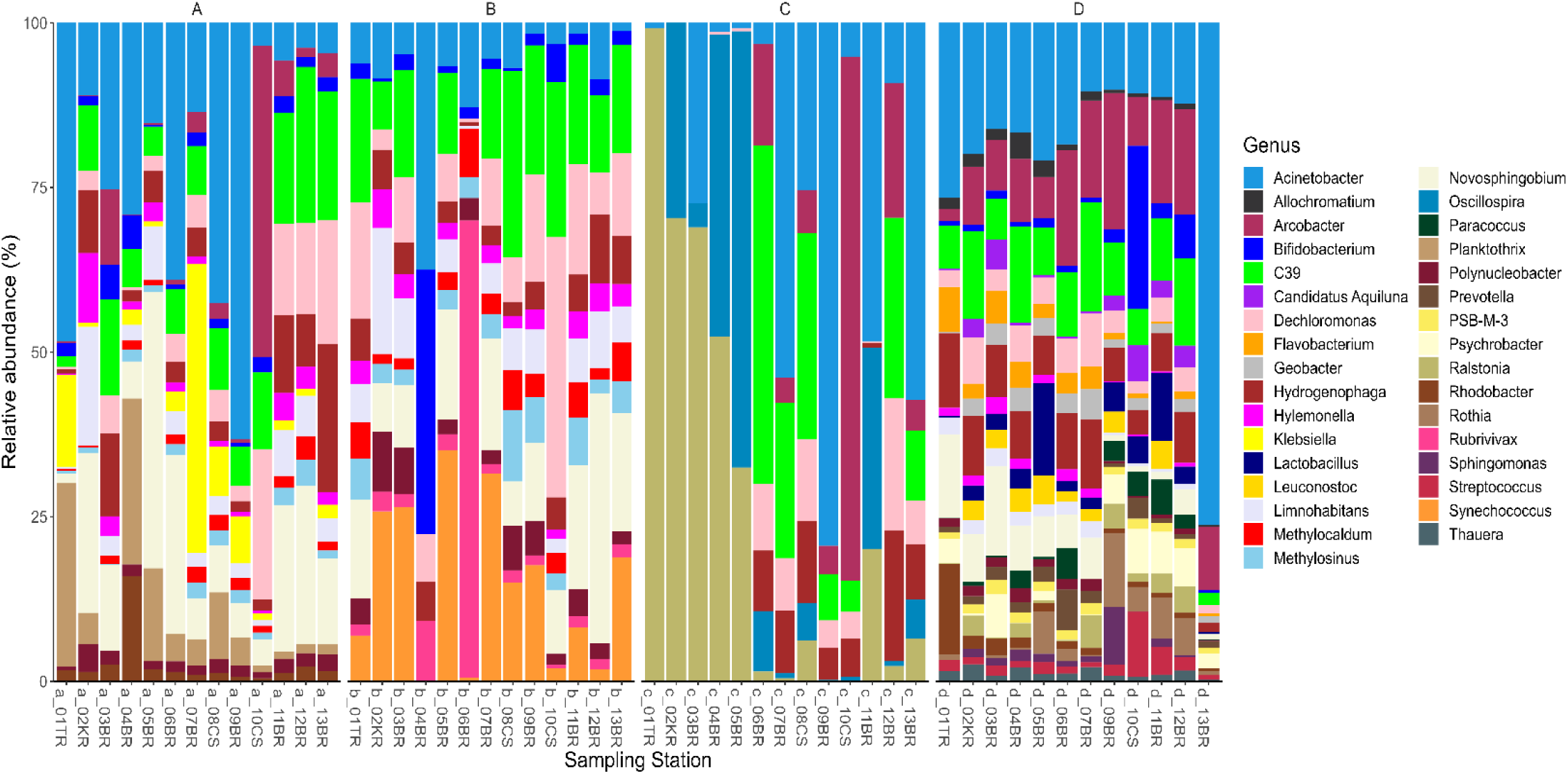
Taxonomic classification of the bacterial community (Genera) at each site (consider relative abundance > 0.5%); A, B, C, and D indicate the pre-monsoon, monsoon, post-monsoon, and winter seasons accordingly.

### 3.3 Temporal Variations of Bacterial Community

#### 3.3.1 At the phylum level

Figure 4 depicted the phylum-level barplot, revealing *Proteobacteria, Actinobacteria,* and *Firmicutes* as major components. *Proteobacteria* emerged as the most abundant bacterial phylum in all water samples, followed by *Actinobacteria, Firmicutes, Bacteroidetes,* and *Cyanobacteria. Proteobacteria* exhibited a gentle decline in relative abundance from pre-monsoon (66.41%) to monsoon (60.40%), followed by an increasing trend from post-monsoon (67.49%) to winter (68.57%). This trend aligns with findings by Dong et al., (2023) at the Bailang River Estuary in China, where *Proteobacteria* constituted the core composition in all their samples. Similar dominant roles of *Proteobacteria* were identified by Liao et al., (2019) and Qiu et al., (2019) in the Maozhou River, China. These bacteria are commonly reported as dominant in contaminated rivers/estuaries (Xiong et al., 2015; Hosen et al., 2017; Guo et al., 2019). *Actinobacteria* displayed a contrasting seasonal pattern compared to *Proteobacteria*, with its relative abundance declining from the monsoon (18.39%) to winter (7.95%). Notably, *Actinobacteria* possesses the capability to decompose organic matter into inorganic forms, facilitating the recycling of molecules by more active microbes, particularly in oligotrophic environments (Wang et al., 2020). *Firmicutes* exhibit greater adaptability to lower temperatures than other bacteria and are proficient in degrading organic nitrogen, suggesting heightened activity in the nitrogen cycle during winter (Wang et al., 2014). In the current study, the abundance of *Firmicutes* experienced a sharp decrease from pre-monsoon (12.09%) to monsoon (7.25%) and then exhibited a significant increase in winter (13.60%). Feng et al., (2022) reported a similar finding in their study at the Bahe River Basin, China, identifying *Firmicutes* as most abundant in disturbed zones, emphasizing the influence of anthropogenic activities. Additionally, *Firmicutes* was found to be the most abundant microbes in a black-odorous section of the Jinchuan River in Nanjing, China, and was proposed as a reliable indicator of fecal pollution (Wu et al., 2019). *Bacteroidetes* exhibited a gradual increase from pre-monsoon (3.65%) to monsoon (4.38%) and the post-monsoon period (10.05%), followed by a decrease in winter (7.20%). Meanwhile, the relative abundance of *Cyanobacteria* doubled abruptly from pre-monsoon (4.35%) to monsoon (9.58%), then swiftly decreased in post-monsoon (2.82%) and maintained this trend in winter (2.69%).

#### 3.3.2 At the genus level

At the genus level (Fig. 5), the six dominant taxa in these samples were *Acinetobacter, C39 (Rhodocyclaceae_*Family*), Arcobacter, Novosphingobium, Dechloromonas,* and *Hydrogenophaga*. However, they exhibited distinct seasonal variations. *Acinetobacter* rapidly decreased from pre-monsoon (21.32%) to monsoon (4.97%), sharply reaching its peak in post-monsoon (34.03%), and then slightly decreasing in winter (22.98%). These bacteria are known for their capacity to oxidize petroleum components through metabolic activities, making them valuable for the biological treatment of oil-contaminated areas (Cai et al., 2021). This genus is commonly found in wastewater and represents a significant component of cultivable microbiota in wastewater and activated sludge. *Acinetobacter* plays a crucial role in nitrification-denitrification processes and phosphorus removal from wastewater (Chen et al., 2019). Additionally, it exhibits resistance to multiple metals, including copper, lead, boron, etc. (Dhakephalkar et al., 1994). On the contrary, *C39 (Rhodocyclaceae_*Family) exhibited a gradual increase from pre-monsoon (11.61%) to monsoon (16.34%), followed by a slow decline in post-monsoon (12.78%) and winter (9.11%). Belonging to the *Proteobacteria* phylum, the genus *C39 (Rhodocyclaceae_*Family) is widely distributed in wastewater remediation systems, playing a crucial role in the removal of various contaminants, including carbon, nitrogen, and phosphorus (Liu et al., 2020). *Arcobacter* mirrored the trend of *C39 (Rhodocyclaceae_*Family), experiencing a swift increase from pre-monsoon (7.65%) to post-monsoon (27.58%) and then a decrease in winter (11.66%). The relative abundance of *Novosphingobium* remained almost the same in pre-monsoon and monsoon seasons, at 15.98% and 16.42%, respectively, before sharply decreasing in winter (5.27%). *Dechloromonas* exhibited a gradual increase from pre-monsoon (9.11%) to monsoon (12.59%), followed by a gradual decrease from post-monsoon (6.55%) to winter (4.10%). In contrast, *Hydrogenophaga* showed a slightly different trend compared to *Dechloromonas*, with a decrease from pre-monsoon (7.18%) to monsoon (5.42%), an increase in post-monsoon (7.48%), and a subsequent decrease in winter (6.87%).

### 3.4 Cluster Analysis

Cluster Analysis (CA) was conducted to clarify the spatial distribution of the research area, as illustrated in the dendrogram produced using Ward’s approach based on the relative abundance of bacteria over the premonsoon, monsoon, postmonsoon, and winter seasons, displayed in Fig. 6 (A and B). The sampling stations at the phylum level exhibit greater variability in the premonsoon, postmonsoon, and winter seasons in terms of their similarities compared to the monsoon period. Conversely, at the genus level, sampling stations during the winter season exhibited greater constraints than in other seasons. This is probably attributable to the persistent pollution levels at the monitoring locations (Alom et al., 2024_Preprint). The heatmap illustrated the relative abundance of microorganisms, with red indicating the highest abundance and green the lowest.

**Fig. 6.**
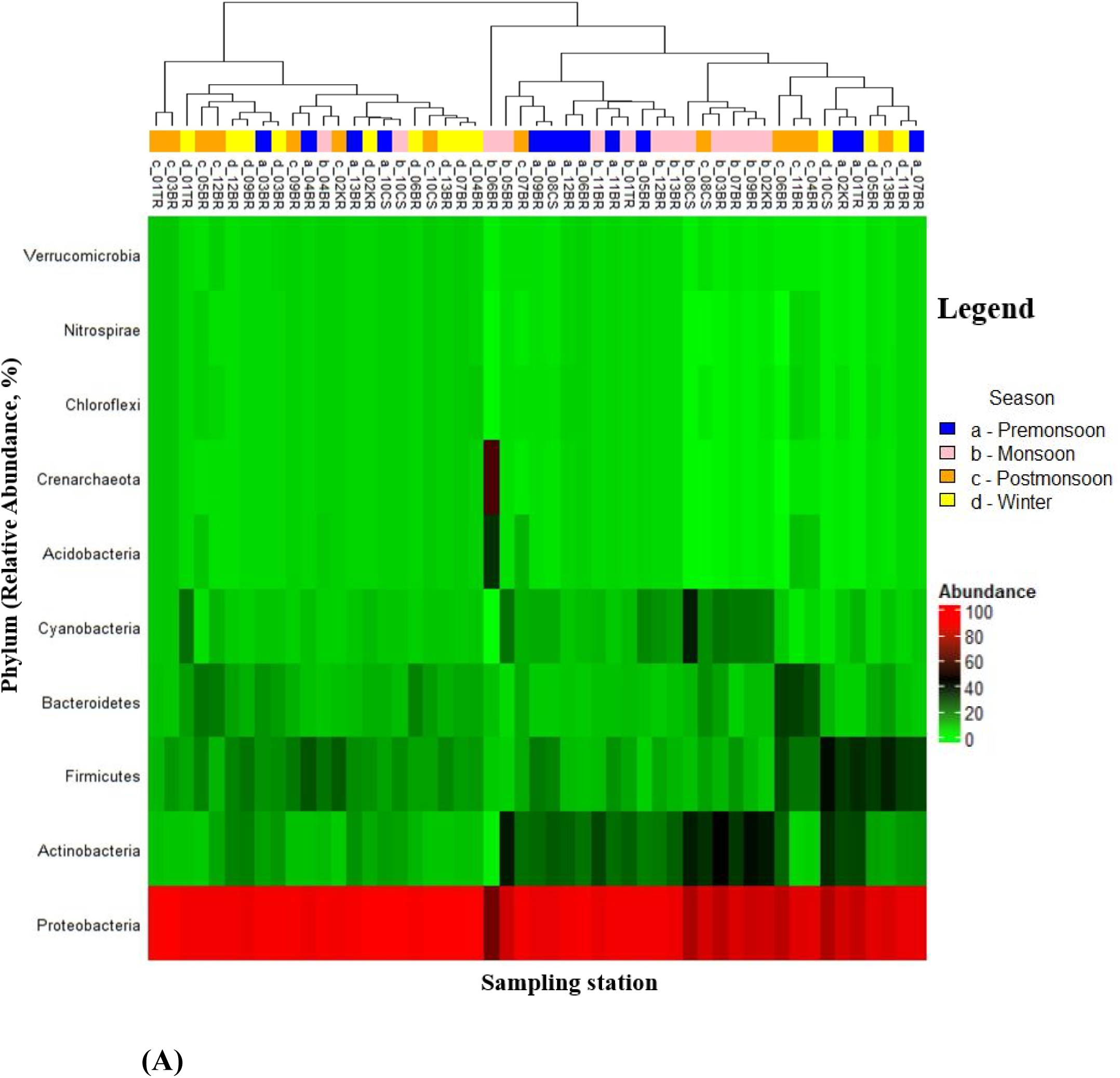
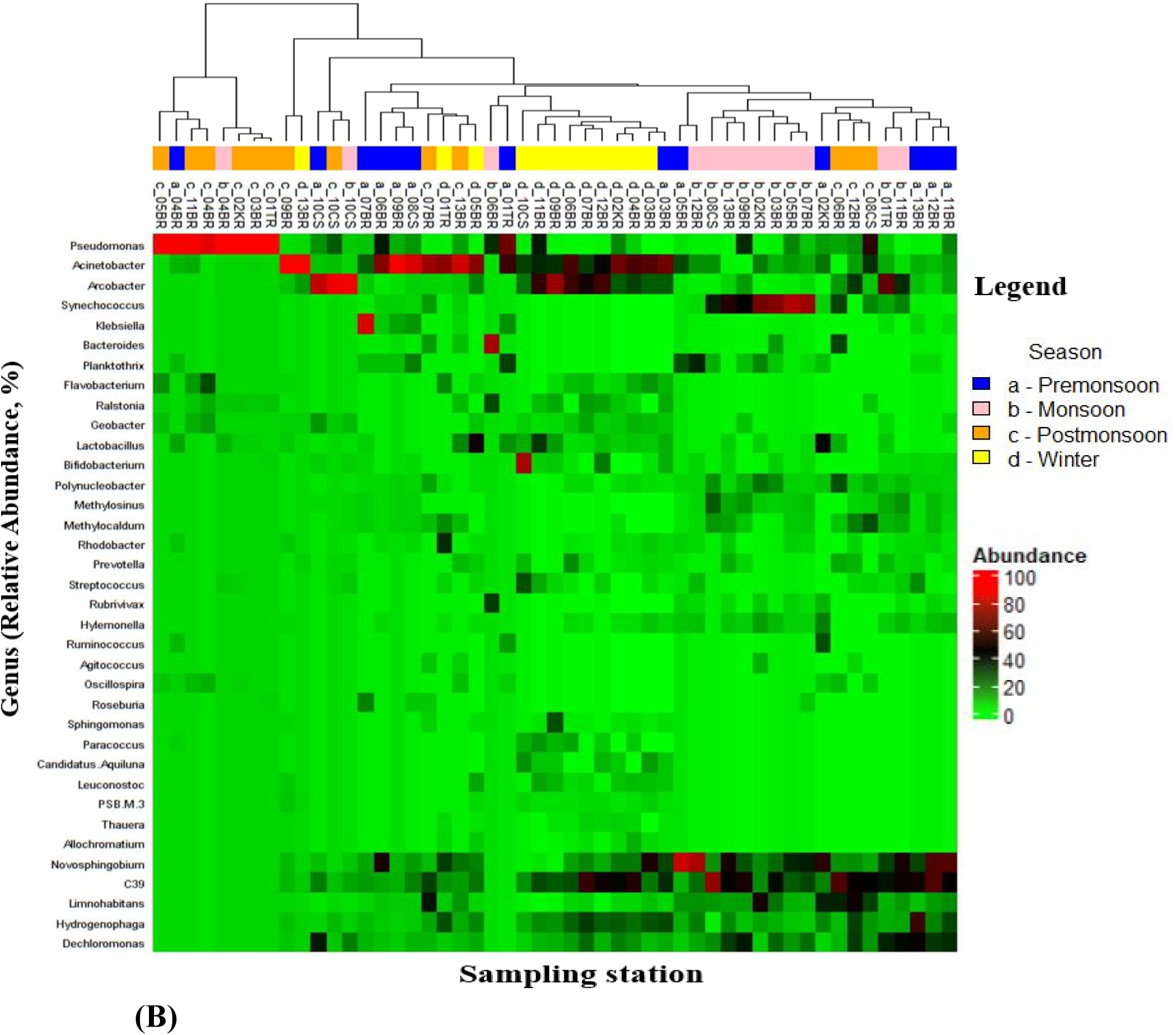
The relative abundance of the bacterial community structure at the **Phylum (A)** and **Genus (B)** levels in different seasons. The dendrogram illustrates the similarities among several stations during the chosen seasons.

### 3.5 Principal Component (PC) Analysis

The combined principal component analysis (Fig. 7A) shows more consistent results in the relative abundance of bacteria at the phylum level across different sampling sites during the winter and pre-monsoon periods, as opposed to the monsoon and post-monsoon periods. Conversely, at the genus level (Fig. 7B), the winter season exhibits relatively stable bacterial abundance across various monitoring stations compared to the other three seasons (pre-monsoon, monsoon, and post-monsoon).

**Fig. 7.**
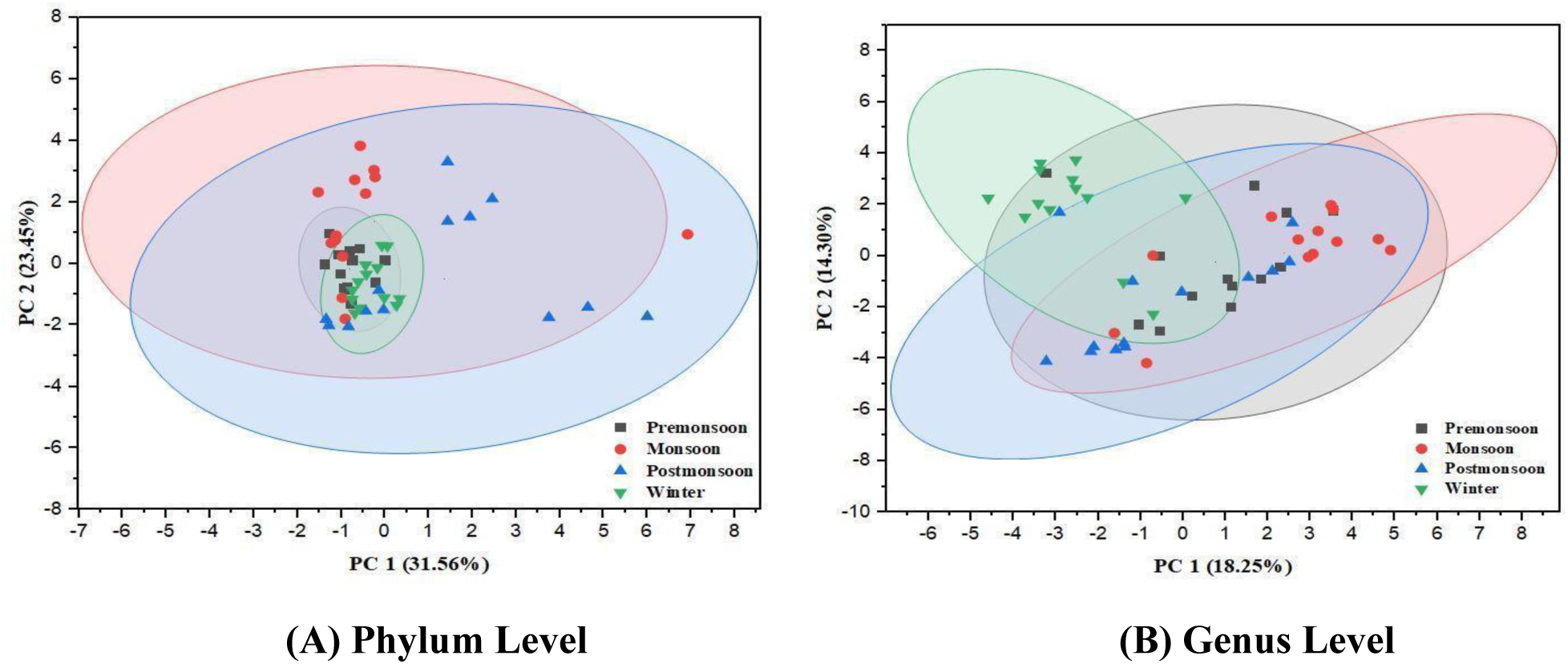
Principal component analysis among the four selected seasons at the **(A)** Phylum and **(B)** Genus level.

These fluctuations in bacterial abundance at different sites and seasons may result from factors such as water dilution and flow patterns, pollution levels at different sites, and water temperature.

### 3.6 Bacterial α- and β-diversity

In Alpha-diversity analysis, both Shannon and Simpson indices yielded consistent results across the four seasons (pre-monsoon, monsoon, post-monsoon, and winter). The analysis highlighted the lowest diversity at sampling site 01TR during the post-monsoon season, while the highest diversity was observed at site 10CS during the winter season (Fig. S2). α-diversity analysis further revealed that the downstream side of the Buriganga River exhibited greater bacterial diversity compared to the upstream side, a result consistent with the relative abundance barplot (downstream > upstream) (Fig. 4 and Fig. 5). This observation may be attributed to various pollution sources along the downstream side of the Buriganga River. From our previous study (Alom et al., 2024_Preprint), based on the results of physicochemical water quality parameters analysis, monitoring site 10CS identified it as the most polluted among the other selected sites. At the junction point of three rivers (Buriganga, Turag, and Karnatali), 03BR exhibited the second-lowest diversity based on Shannon and Simpson indices Figure S2. To assess the variation in seasonal diversity, the Wilcoxon.test (Fig. 8A) was employed to determine the p-value. Due to the minimal changes in water quality parameters (Alom et al., 2024_Preprint) and bacterial richness at the selected sampling sites, the result of α-diversity showed nonsignificant (p > 0.05) across the three seasons (pre-monsoon, monsoon, and post-monsoon). However, a significant difference was observed between winter and any other season, including pre-monsoon, monsoon, and post-monsoon (p < 0.05).

**Fig. 8.**
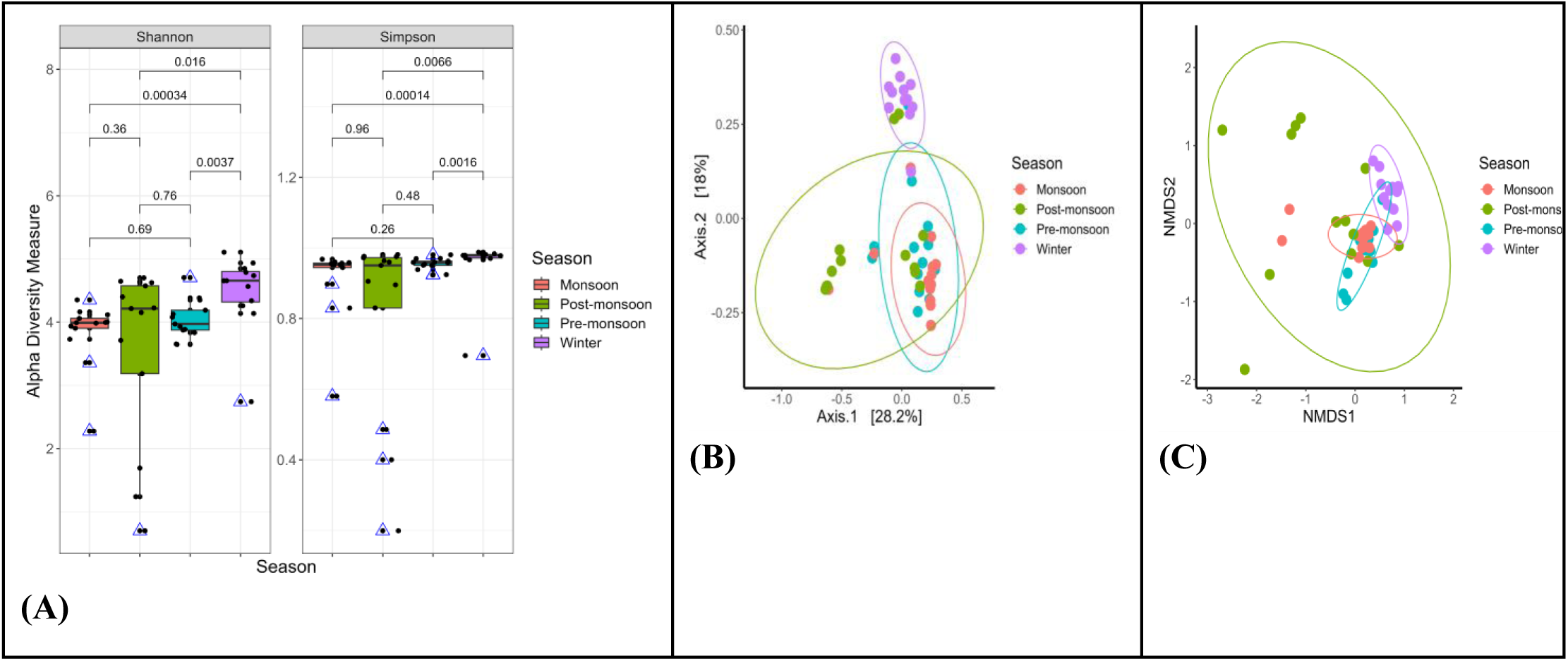
Alpha and Beta diversity. **(A)** Alpha diversity with wilcoxon.test for indicating *P*- value. **(B)** Principal coordinates analysis (PCoA) of the bacterial communities. PC 1 and 2 explained 28.2% and 18% of the variance, respectively and **(C)** Non-metric multidimensional scaling (NMDS).

A Principal Coordinates Analysis (PCoA) of the Bray–Curtis distance, along with a PERMANOVA test was utilized to assess the spatial and temporal distribution of bacterial communities in the water samples. Figure 8B illustrates that the samples were categorized into four clusters, showcasing distinct seasonal changes. Notably, three clusters representing bacterial communities during pre-monsoon, monsoon, and post-monsoon were closely situated, while winter stood alone, displaying a slight distance from the other seasons. This observation was consistent with the results of the Wilcoxon.test, which further highlighted the differences in bacterial communities at the Operational Taxonomic Unit (OTU) level (Fig. 8C).

## 4. Discussion

Various forms of industrial pollution can disrupt the balance of river ecological functions (Ruggiero et al., 2006; Inamdar et al., 2012; Zhang et al., 2021). The bacterial community composition and its shifts were investigated through Illumina MiSeq sequencing at different locations along the Buriganga River, impacted by diverse pollutant sources from various locations. In this study, the predominant bacterial community composition comprised the phyla *Proteobacteria, Actinobacteria,* and *Firmicutes* (Fig. 4). This result is aligned with findings from other studies (Staley et al., 2013; Bai et al., 2014; Ruiz-Gonzalez et al., 2015; Ibekwe et al., 2016).

In our investigation, *Proteobacteria* were the dominant group across all sampling points during the four selected seasons. Known as a key prokaryotic group in aquatic environments (Garrity et al., 2015), *Proteobacteria* encompass taxonomic groups involved in a variety of biogeochemical processes within aquatic ecosystems (Liu et al., 2014; Zhang et al., 2014). Their widespread presence is likely due to their metabolic versatility, adaptability to challenging environments, and rapid growth rates (Spain et al., 2009). Moreover, elevated nutrient levels in the water may be linked to an increase in the relative abundance of *Proteobacteria* (Wu et al., 2022).

*Actinobacteria* emerged as the second most enriched phylum in this study, playing a vital role in the carbon cycle within freshwater ecosystems (Rifaat, 2003). It is reported Total Phosphorus (TP) has an active involvement in the fluctuation of *Actinobacteria*’s abundance in eutrophic environments (Zhang et al., 2021). Additionally, Wang et al., (2018) indicated in their study that TP is a key factor in shaping the microbial community in surface water. Phosphorus is generally acknowledged as a significant driver for biological metabolisms in flowing waters and requires careful management to mitigate eutrophication impacts resulting from urbanization (Withers and Jarvie, 2008). *Actinobacteria* solubilizes the insoluble phosphates in river sediments, making the phosphorus available for uptake by plants and other organisms in the aquatic ecosystem and playing an important role in the phosphorus cycle (Dastager & Damare, 2013).

*Firmicutes,* another phylum, exhibited maximum abundance during winter at sampling station 10CS, located inside the canal and previously identified as the most polluted site in our prior study (Alom et al., 2024_Preprint). This heightened presence may be attributed to the substantial enrichment of untreated domestic and industrial sewage pollutants. Excessive organic load might have created anaerobic conditions that facilitated fermentation and subsequently low pH (Ballester et al., 1999). *Firmicutes* received a competitive edge here because they can degrade chemically complex organics, can withstand high toxicity and pH, and comprise mostly anaerobic and facultative anaerobic bacteria (Filippidou et al., 2016). The depletion in number during the monsoon and increase during winter can also be explained by the fact that high water flow during monsoon might have reduced toxicity and increased oxygen (Hassan et al., 2023). *Firmicutes* are recognized as an indicator of fecal contamination and a marker for identifying human feces in water by Zheng et al., (2009). Consequently, it can be inferred that untreated sewage effluents influence both the physicochemical properties of river water and the composition of bacterial communities. These observations are in line with findings in other studies, highlighting microbial community alterations due to wastewater effluents (Drury et al., 2013; Garcia-Armisen et al., 2014).

*Cyanobacteria* grow rapidly in nutrient-rich water bodies, and actively participate in different geochemical cycles and photosynthesis, but may cause eutrophication. It requires a specific amount of light for photosynthesis and is sensitive to temperature (Muhetaer et al., 2020). Less number of cyanobacteria during winter might indicate the effect of less availability of light and cold temperatures (Lürling et al., 2017).

*Bacteroidetes* was another predominant phylum in our observations, potentially originating from untreated wastewater in residential areas containing pollutants such as ammonia and feces. Wéry et al., (2008) proposed certain species within this phylum as effective alternative fecal indicators, and Liu et al., (2008) identified *Bacteroidetes* as a denitrifying agent.

In Genus-level, analysis across the four seasons showed increased bacterial diversity in winter. This finding aligns with evidence that colder temperatures, reduced oxygen availability, slower water flow, and high nutrient concentrations can suppress autochthonous bacterial activity (Saifullah et al., 2013). These conditions may have allowed allochthonous bacteria to enter, enhancing the observed diversity in the winter season (Luo et al., 2020; Barcina et al., 1997).

In our observation, *Acinetobacter* exhibited the highest abundance throughout the year in the Buriganga River, particularly during the post-monsoon and winter seasons when heavy metal concentrations were elevated (Alom et al., 2024_Preprint). Some studies have suggested that wastewater containing heavy metals from iron and steel plants can decompose soil, releasing heavy metals into rivers through acidic runoff (Holding, 2004). This process may contribute significantly to the extensive enrichment of *Acinetobacter*. The abundance of this genus can serve as a sensitive indicator of heavy metal content in rivers (Zhang et al., 2021).

The high abundance of *Pseudomonas* and *Ralstonia* at stations 01TR, 02KR, and 03BR—located outside the main flow of the Buriganga River—during the post-monsoon season suggested potential antibiotic contamination as these genera are well known for their strong resistance to multiple antibiotics and their ability to thrive in environments contaminated with antibiotics (Luczkiewicz et al., 2015; Ryan & Adley, 2013). Their dominance indicated the likely presence of antibiotic residues in these areas, promoting the survival and growth of resistant bacteria.

*Acinetobacter,* which is well-known for their antibiotic resistance and are often found in hospital wastewater and sewage (Moubareck & Halat, 2020). Their prevalence at multiple sites, especially during winter and post-monsoon periods, indicates possible antibiotic contamination in these water bodies from nearby hospitals, which might have lead to the selection of antibiotic-resistant bacteria. Moreover, *Acinetobacter’s* high tolerance to heavy metals and its ability to detoxify them provided an advantage over *Pseudomonas* and *Ralstonia* in the Buriganga River which showed high levels of heavy metal contamination (Ghaima et al., 2018; Alom et al., 2024_Preprint)

The presence of specific bacteria can indicate the presence of certain heavy metals. For instance, *Geobacter* is known for its ability to reduce and detoxify metals like iron and uranium (Golchereh Mirlahiji & Eisazadeh, 2014). Its presence suggests that there may be heavy metal pollution, especially with metals that *Geobacter* can process. *Dechloromonas*, found at station 10CS, can reduce chlorinated compounds and tolerate heavy metals, indicating potential contamination with both chlorinated solvents and heavy metals.

*Arcobacter* species are emerging pathogens known to cause gastroenteritis in humans (Kayman et al., 2012), and their prevalence at site 10 CS suggests an elevated health risk in that area. Additionally, the widespread presence of *Acinetobacter* and *Pseudomonas* throughout the river signifies a potential health hazard associated with using the water from the Buriganga.

This current study observed a significant association between Zn, TP, BOD5 and COD with the diversity of microbial communities along the Buriganga River (Alom et al., 2024_Preprint). During the winter season, these parameters exhibited peak values, coinciding with the maximum bacterial diversity. In contrast, the post-monsoon season displayed the lowest microbial diversity alongside minimum values for physicochemical parameters (Zn, TP, BOD5, and COD). Numerous previous studies have illustrated the impact of metal selection on microbial communities in river ecosystems (Feris et al., 2004; Morin et al., 2008; Yin et al., 2015). Wang et al., (2018) discovered in their investigation along the Jialing River in China that Zinc (Zn) and Iron (Fe) were primarily linked to the spatial distribution of microbial communities in urban rivers. Our exploration revealed lower microbial diversity and OTU richness on the upstream side (01TR, 02KR, 03BR, 04BR, 05BR, and 06BR) compared to the downstream side of the river. This discrepancy could be happening due to concentration of pollutants and the substantial enrichment of species well-adapted to domestic and industrial pollution conditions. The results suggest that the composition and diversity of bacterial communities within the same river is influenced by various environmental factors, and distinct types of industrial and residential pollutions.

## 5. Conclusion

This study employed 16S rRNA gene-based metagenomics sequencing to examine the spatial and seasonal diversity of bacterial communities in the Buriganga River, considering various inputs from residential and industrial pollution sources. Our findings underscored that the bacterial community in the river is intricately shaped by diverse environmental factors, with distinct types of pollution exerting varied effects on the richness and evenness of the bacterial community within the same river. While the bacterial diversity did not significantly differ among the three seasons—pre-monsoon, monsoon, and post-monsoon—a significant disparity was observed between winter and any other selected season. This observation aligns with the heightened concentration of physicochemical water parameters in the winter season, indicating elevated pollution levels compared to other seasons. Our analyses highlighted Zn, TP, BOD, and COD as the main but not the only factors influencing the community structure of bacteria, impacting the distribution of dominant phyla and genera. Additionally, the identification of potential pathogens such as *Arcobacter, Acinetobacter,* and *Pseudomonas* in the samples raises concerns about potential risks to human health. The outcomes of this research not only offer insights for monitoring and managing the Buriganga River ecosystem but also serve as a valuable data bank for future investigations, aiding researchers in comprehending the microbial ecology of urban river water.

## References

1. Alom, M. M., Sultana, M & Rahman, S. M., (2024). Spatial and temporal variation of water quality of Buriganga River, Dhaka. Available at SSRN: https://ssrn.com/abstract=4890564 or 10.2139/ssrn.4890564 (Preprint).

2. Bai Y, Qi W, Liang J, Qu J. (2014). Using high-throughput sequencing to assess the impacts of treated and untreated wastewater discharge on prokaryotic communities in an urban river. Appl Microbiol Biotechnol. 98(4):1841–1851. DOI: 10.1007/s00253-013-5116-2

3. Bååth, E., & Kritzberg, E. S. (2024). Temperature Adaptation of Aquatic Bacterial Community Growth Is Faster in Response to Rising than to Falling Temperature. Microbial Ecology, 87(1), 1–8. 10.1007/s00248-024-02353-8

4. Ballester, M. V., Martinelli, L. A., Krusche, A. V., Victoria, R. L., Bernardes, M., & Camargo, P. B. (1999). Effects of increasing organic matter loading on the dissolved O2, free dissolved CO2 and respiration rates in the Piracicaba River basin, Southeast Brazil. Water Research, 33(9), 2119–2129. 10.1016/S0043-1354(98)00438-2

5. Barcina, I., Lebaron, P., & Vives-Rego, J. (1997). Survival of allochthonous bacteria in aquatic systems: a biological approach. FEMS Microbiology Ecology, 23(1), 1–9. 10.1111/J.1574-6941.1997.TB00385.X

6. Bhuiyan, M. A. H., Dampare, S. B., Islam, M. A., & Suzuki, S. (2015). Source apportionment and pollution evaluation of heavy metals in water and sediments of Buriganga River, Bangladesh, using multivariate analysis and pollution evaluation indices. Environmental monitoring and assessment, 187, 1–21. DOI: 10.1007/s10661-014-4075-0

7. Breton-Deval L, Sanchez-Flores A, Juarez K, Vera-Estrella R. (2019). Integrative study of microbial community dynamics and water quality along the Apatlaco river. Environ Pollut. 255(Pt 1):113158. 10.1016/j.envpol.2019.113158

8. Cai, Y., Wang, R., Rao, P., Wu, B., Yan, L., Hu, L., Park, S., Ryu, M., & Zhou, X. (2021). Bioremediation of petroleum hydrocarbons using acinetobacter sp. SCYY-5 isolated from contaminated oil sludge: Strategy and effectiveness study. International Journal of Environmental Research and Public Health, 18(2), 1–14. 10.3390/ijerph18020819

9. Callahan, B. J., McMurdie, P. J., Rosen, M. J., Han, A. W., Johnson, A. J. A., and Holmes, S. P. (2016). DADA2: high resolution sample inference from illumina amplicon data. Nat. Methods 13, 581–583. doi: 10.1038/nmeth. 3869

10. Caporaso, J.G., Kuczynski, J., Stombaugh, J., Bittinger, K., Bushman, F.D., Costello, E.K., (2010b). QIIME allows analysis of high-throughput community sequencing data. Nat. Methods 7, 335–336. doi:10.1038/nmeth.f.303

11. Chen, K., Jiao, J.J., Huang, J., Huang, R. (2007). Multivariate statistical evaluation of trace elements in groundwater in a coastal area in Shenzhen, China. Environ. Pollut. 147 (3), 771–780. 10.1016/j.envpol.2006.09.002

12. Chen, L., Tsui, M.M.P., Lam, J.C.W., Hu, C., Wang, Q., Zhou, B., Lam, P.K.S. (2019). Variation in microbial community structure in surface seawater from Pearl River Delta: Discerning the influencing factors. Sci. Total Environ. 660, 136–144. 10.1016/j.scitotenv.2018.12.480.

13. Chen, Y., Liu, Y., Li, Y., Wu, Y., Chen, Y., Zeng, G., Zhang, J., Li, H. (2017). Influence of biochar on heavy metals and microbial community during composting of river sediment with agricultural wastes. Bioresour. Technol. 243, 347–355. https://doi.org/10.1016/j.biortech.2017.06.100.

14. Dastager, S. G., & Damare, S. (2013). Marine Actinobacteria showing Phosphate solubilizing efficiency in Chorao Island, Goa, India. Curr. Microbiol, 66(5), 421–427. http://www.ncbi.nlm.nih.nov

15. Dhakephalkar, P. K., & Chopade, B. A. (1994). High levels of multiple metal resistance and its correlation to antibiotic resistance in environmental isolates of Acinetobacter. Biometals, 7(1), 67–74. doi:10.1007/bf00205197

16. DNeasy PowerWater Kit Handbook (2022). QIAGEN, USA. https://www.qiagen.com/us/resources/resourcedetail?id=75765ef9-2a6f-4f5d-a36b-dbd9beb43079&lang=en&srsltid=AfmBOopQtRmpe_N3zqt28zTcbWCYxffW4DPbMx9VMPnCWyjXay-pkF4v (accessed 10 August 2024)

17. Dong, W., Cui, Z., Zhao, M., & Li, J. (2023). Seasonal and Spatial Variations of Bacterial Community Structure in the Bailang River Estuary. Journal of Marine Science and Engineering, 11(4), 825. 10.3390/jmse11040825

18. Douglas, G. M., Maffei, V. J., Zaneveld, J. R., Yurgel, S. N., Brown, J. R., Taylor, C. M., et al. (2020). PICRUSt2 for prediction of metagenome functions. Nat. Biotechnol. 38, 685–688. doi: 10.1038/s41587-020-0548-6

19. Drury, B., Rosi-Marshall, E., Kelly, J.J. (2013). Wastewater treatment effluent reduces the abundance and diversity of benthic bacterial communities in urban and suburban rivers. Appl. Environ. Microbiol. 79, 1897–1905. 10.1128/AEM.03527-12

20. Fatema, K., Begum, M., Zahid, M., & Hossain, M. E. (2018). Water quality assessment of the river Buriganga. Bangladesh. Journal of Biodiversity Conservation and Bioresource Management, 4(1), 47–54. 10.3329/jbcbm.v4i1.37876

21. Feng, W., Gao, J., Wei, Y., Liu, D., Yang, F., Zhang, Q., & Bai, Y. (2022). Pattern changes of microbial communities in urban rivers affected by anthropogenic activities and their environmental driving mechanisms. Environmental Sciences Europe, 34(1), 1–13. 10.1186/s12302-022-00669-1

22. Feris, K.P., Ramsey, P.W., Frazar, C., Rillig, M., Moore, J.N., Gannon, J.E., et al. (2004). Seasonal dynamics of shallow-hyporheic-zone microbial community structure along a heavy metal contamination gradient. Appl. Environ. Microbiol. 70, 2323–2331. 10.1128/AEM.70.4.2323-2331.2004

23. Filippidou, S., Wunderlin, T., Junier, T., Jeanneret, N., Dorador, C., Molina, V., Johnson, D. R., & Junier, P. (2016). A combination of extreme environmental conditions favor the prevalence of endospore-forming firmicutes. Frontiers in Microbiology, 7(NOV), 223369. 10.3389/fmicb.2016.01707

24. Gao, Y., Zhang, W., Li, Y., Wu, H., Yang, N., Hui, C. (2021). Dams shift microbial community assembly and imprint nitrogen transformation along the Yangtze River. Water Res. 189, 116579 10.1016/j.watres.2020.116579.

25. Garcia-Armisen, T., Inceoglu, O., Ouattara, N.K., Anzil, A., Verbanck, M.A., Brion, N., et al. (2014). Seasonal variations and resilience of bacterial communities in a sewage polluted urban river. PLoS One 9, e92579. 10.1371/journal.pone.0092579

26. Garcia-Mazcorro J, Treviño-Espinosa R, Medina-Ponce M, MarroquinCardona A, Cruz-Valdéz J, Ramírez-Martínez C. (2016). Massive molecular description of microorganisms in an urban river. Toxicol Lett 259:S127. 10.1016/j.toxlet.2016.07.326

27. Garrity GM, Bell JA, Lilburn T. (2015). Proteobacteria phyl. nov. Bergey’s Manual of Systematics of Archaea and Bacteria. John Wiley & Sons, Ltd. 10.1002/9781118960608.pbm00022

28. Ghaima, K. K., Lateef, N. S., & Khaz’al, Z. T. (2018). Heavy metal and antibiotic resistance of Acinetobacter spp. isolated from diesel fuel polluted soil. Journal of Advanced Laboratory Research in Biology, 9(2), 58–64.

29. Golchereh Mirlahiji, S., & Eisazadeh, K. (2014). Bioremediation of Uranium Via Geobacter Spp. Journal of Research and Development, 1(12), 52–58.

30. Guo, X., Yang, Y., Niu, Z., Lu, D., Zhu, C., Feng, J., et al., (2019). Characteristics of microbial community indicate anthropogenic impact on the sediments along the Yangtze Estuary and its coastal area, China. Sci. Total Environ. 648, 306e314. 10.1016/j.scitotenv.2018.08.162

31. Hassan, H. Bin, Moniruzzaman, M., Majumder, R. K., Ahmed, F., Quaiyum Bhuiyan, M. A., Ahsan, M. A., & Al-Asad, H. (2023). Impacts of seasonal variations and wastewater discharge on river quality and associated human health risks: A case of northwest Dhaka, Bangladesh. Heliyon, 9(7). 10.1016/J.HELIYON.2023.E18171

32. Holding B. (2004). Water treatment & air purification. Rotterdamseweg: Lenntech; p. 58–78.

33. Hosen, J.D., Febria, C.M., Crump, B.C., Palmer, M.A., (2017). Watershed urbanization is linked to differences in stream bacterial community composition. Front. Microbiol. 8, 01452. 10.3389/fmicb.2017.01452

34. Huang, F., Zou, S., Deng, D., Lang, H., and Liu, F. (2019). Antibiotics in a typical karst river system in China: Spatiotemporal variation and environmental risks. Sci. Total Environ. 650, 1348–1355. doi: 10.1016/j.scitotenv.2018.09.131

35. Ibekwe AM, Ma J, Murinda SE. (2016). Bacterial community composition and structure in an Urban River impacted by different pollutant sources. Sci Total Environ. 566–567:1176–1185. 10.1016/j.scitotenv.2016.05.168

36. Inamdar S, Finger N, Singh S, Mitchell M, Levia D, Bais H, Scott D, McHale P. (2012). Dissolved organic matter (DOM) concentration and quality in a forested mid-Atlantic watershed, USA. Biogeochemistry. 108(1–3):55–76. DOI 10.1007/s10533-011-9572-4

37. Islam, M. M., Rahman, S. L., Ahmed, S. U., & Haque, M. K. I. (2014). Biochemical characteristics and accumulation of heavy metals in fishes, water and sediments of the river Buriganga and Shitalakhya of Bangladesh. Journal of Asian scientific research, 4(6), 270.

38. Islam, S. M. D., & Azam, G. (2015). Seasonal variation of physicochemical and toxic properties in three major rivers; Shitalakhya, Buriganga and Turag around Dhaka city. Bangladesh. J. Biodivers. Environ. Sci, 7(3), 120–131.

39. Kayman, T., Abay, S., Hizlisoy, H., Ibrahim Atabay, H., Serdar Diker, K., & Aydin, F. (2012). Emerging pathogen Arcobacter spp. in acute gastroenteritis: Molecular identification, antibiotic susceptibilities and genotyping of the isolated arcobacters. Journal of Medical Microbiology, 61(PART 10), 1439–1444. 10.1099/jmm.0.044594-0

40. Kostanjsek, R., Lapanje, A., Drobne, D., Nikcevic, S., Perović, A., Zidar, P., et al., (2005). Bacterial community structure analyses to assess pollution of water and sediments in the Lake Shkodra/Skadar, Balkan Peninsula (8 pp). Environ. Sci. Pollut. Res. Int. 12, 361–368.

41. Li, J., Chen, Q., Li, Q., Zhao, C., Feng, Y. (2021a). Influence of plants and environmental variables on the diversity of soil microbial communities in the Yellow River Delta Wetland. China. Chemosphere 274, 129967. 10.1016/j.chemosphere.2021.129967.

42. Li, Y., Gao, Y., Zhang, W., Wang, C., Wang, P., Niu, L., et al. (2019). Homogeneous selection dominates the microbial community assembly in the sediment of the Three Gorges Reservoir. Sci. Total Environ. 690, 50e60. 10.1016/j.scitotenv.2019.07.014

43. Liang, Z., Fang, W., Luo, Y., Lu, Q., Juneau, P., He, Z., Wang, S. (2021). Mechanistic insights into organic carbon-driven water blackening and odorization of urban rivers. J. Hazard. Mater. 405, 124663 10.1016/j.jhazmat.2020.124663.

44. Liang Z, Siegert M, Fang W, Sun Y, Jiang F, Lu H, Chen G-H, Wang S. (2018). Blackening and odorization of urban rivers: a bio-geochemical process. FEMS Microbiol Ecol. 94(3). 10.1093/femsec/ fix180

45. Liao, H., Yu, K., Duan, Y., Ning, Z., Li, B., He, L., et al. (2019). Profiling microbial communities in a watershed undergoing intensive anthropogenic activities. Sci. Total Environ. 647, 1137e1147. 10.1016/j.scitotenv.2018.08.103

46. Lindström, E.S., Kamst-Van Agterveld, M.P., Zwart, G. (2005). Distribution of typical freshwater bacterial groups is associated with pH, temperature, and lake water retention time. Appl. Environ. Microbiol. 71, 8201–8206. DOI: 10.1128/AEM.71.12.8201-8206.2005

47. Liu J, Liu X, Wang M, Qiao Y, Zheng Y, Zhang X (2015). Bacterial and archaeal communities in sediments of the north Chinese marginal seas. Microb Ecol 70(1):105–117. 10.1007/s00248-014-0553-8

48. Liu, S., Daigger, G.T., Liu, B., Zhao, W., Liu, J., (2020). Enhanced performance of simultaneous carbon, nitrogen and phosphorus removal from municipal wastewater in an anaerobic-aerobic-anoxic sequencing batch reactor (AOA-SBR) system by alternating the cycle times. Bioresour. Technol. 301, 122750 10.1016/j.biortech.2020.122750.

49. Liu X, Gao C, Zhang A, Jin P, Wang L, Feng L. (2008). The nos gene cluster from gram-positive bacterium Geobacillus thermodenitrificans NG80-2 and functional characterization of the recombinant NosZ. FEMS Microbiol Lett. 289(1):46–52. 10.1111/j.1574-6968.2008.01362.x

50. Liu Y, Zhang J, Zhao L, Zhang X, Xie S. (2014). Spatial distribution of bacterial communities in high altitude freshwater wetland sediment. Limnology. 15(3):249–256. DOI 10.1007/s10201-014-0429-0

51. Luczkiewicz, A., Kotlarska, E., Artichowicz, W., Tarasewicz, K., & Fudala-Ksiazek, S. (2015). Antimicrobial resistance of Pseudomonas spp. isolated from wastewater and wastewater-impacted marine coastal zone. Environmental Science and Pollution Research, 22(24), 19823–19834. 10.1007/s11356-015-5098-y

52. Lundgaard, A.S.B., Treusch, A.H., Stief, P., Thamdrup, B., Glud, R.N., (2017). Nitrogen cycling and bacterial community structure of sinking and aging diatom aggregates. Aquat. Microb. Ecol. 79, 85–99. 10.3354/ame01821

53. Luo, X., Xiang, X., Yang, Y., Huang, G., Fu, K., Che, R., & Chen, L. (2020). Seasonal effects of river flow on microbial community coalescence and diversity in a riverine network. FEMS Microbiology Ecology, 96(8). 10.1093/FEMSEC/FIAA132

54. Lürling, M., Van Oosterhout, F., & Faassen, E. (2017). Eutrophication and warming boost cyanobacterial biomass and microcystins. Toxins, 9(2), 1–16. 10.3390/toxins9020064

55. Mark Ibekwe, A., Leddy, M.B., Bold, R.M., Graves, A.K., (2012). Bacterial community composition in low-flowing river water with different sources of pollutants. FEMS Microbiol. Ecol. 79, 155–166. 10.1111/j.1574-6941.2011.01205.x

56. McDonald, D., Price, M. N., Goodrich, J., Nawrocki, E. P., DeSantis, T. Z., Probst, A., … & Hugenholtz, P. (2012). An improved Greengenes taxonomy with explicit ranks for ecological and evolutionary analyses of bacteria and archaea. The ISME journal, 6(3), 610–618. 10.1038/ismej.2011.139

57. Morin, S., Duong, T., Dabrin, A., Coynel, A., Herlory, O., Baudrimont, M., et al. (2008). Long-term survey of heavy-metal pollution, biofilm contamination and diatom community structure in the Riou Mort watershed, South-West France. J. Environ. Pollut. 151, 532–542. 10.1016/j.envpol.2007.04.023

58. Moubareck, C. A., & Halat, D. H. (2020). Insights into Acinetobacter baumannii: A Review of Microbiological, Virulence, and Resistance Traits in a Threatening Nosocomial Pathogen. Antibiotics (Basel), 9(3), 1–29. 10.3390/antibiotics9030119

59. Muhetaer, G., Asaeda, T., Jayasanka, S. M. D. H., Baniya, M. B., Abeynayaka, H. D. L., Rashid, M. H., & Yan, H. Y. (2020). Effects of Light Intensity and Exposure Period on the Growth and Stress Responses of Two Cyanobacteria Species: Pseudanabaena galeata and Microcystis aeruginosa. Water 2020, Vol. 12, Page 407, 12(2), 407. 10.3390/W12020407

60. Murphy CL, Biggerstaf J, Eichhorn A, Ewing E, Shahan R, Soriano D, Stewart S, Vanmol K, Walker R, Walters P, Elshahed M, Youssef N (2021). Genomic characterization of three novel Desulfobacterota classes expand the metabolic and phylogenetic diversity of the phylum. Environ Microbiol 23(8):4326–4343. 10.1111/1462-2920.15614

61. Ouyang, L., Chen, H., Liu, X., Wong, M. H., Xu, F., Yang, X., … & Li, S. (2020). Characteristics of spatial and seasonal bacterial community structures in a river under anthropogenic disturbances. Environmental Pollution, 264, 114818. 10.1016/j.envpol.2020.114818

62. Peterson, D., Bonham, K. S., Rowland, S., Pattanayak, C. W., Resonance Consortium, & Klepac-Ceraj, V. (2021). Comparative analysis of 16S rRNA gene and metagenome sequencing in pediatric gut microbiomes. Frontiers in microbiology, 12, 670336. 10.3389/fmicb.2021.670336

63. Qin, W., Han, D., Song, X., and Liu, S. (2021). Sources and migration of heavy metals in a karst water system under the threats of an abandoned Pb–Zn mine. Southwest China. Environ. Pollut. 277:116774. doi: 10.1016/j.envpol.2021.116774

64. Qin, Y., Hao, F., Zhang, D., Lang, Y., and Wang, F. (2020). Accumulation of organic carbon in a large canyon reservoir in Karstic area, Southwest China. Environ. Sci. Pollut. Res. 27, 25163–25172. doi: 10.1007/s11356-020-08724-1

65. Qiu, W., Sun, J., Fang, M., Luo, S., Tian, Y., Dong, P., et al., (2019). Occurrence of antibiotics in the main rivers of Shenzhen, China: association with antibiotic resistance genes and microbial community. Sci. Total Environ. 653, 334e341. 10.1016/j.scitotenv.2018.10.398

66. Raat AJ. (2001). Ecological rehabilitation of the Dutch part of the River Rhine with special attention to the fish. Regul Rivers: Res Mgmt. 17(2):131–144. 10.1002/rrr.608

67. Rahman, A., & Al Bakri, D. (2010). A study on selected water quality parameters along the River Buriganga, Bangladesh. Iranian (Iranica) Journal of Energy & Environment, 1(2).

68. Rajaram T, Das A. (2008). Water pollution by industrial effluents in India: discharge scenarios and case for participatory ecosystem specific local regulation. Futures. 40(1):56–69. 10.1016/j.futures.2007.06.002

69. Rifaat HM. (2003). The biodiversity of actinomycetes in the River Nile exhibiting antifungal activity. J Mediterranean Ecol. 4:5–8.

70. Ruggiero A, Solimini AG, Carchini G. (2006). Effects of a wastewater treatment plant on organic matter dynamics and ecosystem functioning in a Mediterranean stream. Ann Limnol Int J Lim. 42(2):97–107. 10.1051/limn/2006014

71. Ruiz-Gonzalez C, Nino-Garc ∼ ıa JP, del Giorgio PA. 2015. Terrestrial origin of bacterial communities in complex boreal freshwater networks. Ecol Lett. 18(11):1198–1206. 10.1111/ele.12499

72. Saifullah, A., Kabir, M., Khatun, A., Roy, S. and Sheikh, M. (2013). Investigation of Some Water Quality Parameters of the Buriganga River. Journal of Environmental Science and Natural Resources, 5(2): 47–52.

73. Ryan, M. P., & Adley, C. C. (2013). The antibiotic susceptibility of water-based bacteria Ralstonia pickettii and Ralstonia insidiosa. Journal of Medical Microbiology, 62(PART7), 1025–1031. 10.1099/jmm.0.054759-0

74. Saifullah, A., Kabir, M., Khatun, A., Roy, S., & Sheikh, M. (2013). Investigation of Some Water Quality Parameters of the Buriganga River. Journal of Environmental Science and Natural Resources, 5(2), 47–52. 10.3329/jesnr.v5i2.14600

75. Sheng Y, Qu Y, Ding C, Sun Q, Mortimer RJ. (2013). A combined application of different engineering and biological techniques to remediate a heavily polluted river. Ecol Eng. 57:1–7. 10.1016/j.ecoleng.2013.04.004

76. Shi Y, Su C, Wang M, Liu X, Ma W (2020). Modern climate and soil properties explain functional structure better than phylogenetic structure of plant communities innorthern China. Front Ecol Evol 8:531947. 10.3389/fevo.2020.531947

77. Sierra, M. A., Li, Q., Pushalkar, S., Paul, B., Sandoval, T. A., Kamer, A. R., … & Saxena, D. (2020). The influences of bioinformatics tools and reference databases in analyzing the human oral microbial community. Genes, 11(8), 878. 10.3390/genes11080878

78. Spain, A. M., Krumholz, L. R., & Elshahed, M. S. (2009). Abundance, composition, diversity and novelty of soil Proteobacteria. ISME Journal, 3(8), 992–1000. 10.1038/ISMEJ.2009.43

79. Sreevidya, M., Gopalakrishnan, S., Kudapa, H., & Varshney, R. K. (2016). Exploring plant growth-promotion actinomycetes from vermicompost and rhizosphere soil for yield enhancement in chickpea. Brazilian Journal of Microbiology, 47(1), 85–95. 10.1016/j.bjm.2015.11.030

80. Staley C, Unno T, Gould TJ, Jarvis B, Phillips J, Cotner JB, Sadowsky MJ. 2013. Application of Illumina next-generation sequencing to characterize the bacterial community of the Upper Mississippi River. J Appl Microbiol. 115(5):1147–1158. 10.1111/jam.12323

81. Sun, Y., Wang, S., Niu, J. (2018). Microbial community evolution of black and stinking rivers during in situ remediation through micro-nano bubble and submerged resin floating bed technology. Bioresour. Technol. 258, 187–194. 10.1016/j.biortech.2018.03.008.

82. Su Z, Dai T, Tang Y, Tao Y, Huang B, Mu Q, Wen D. (2018) Sediment bacterial community structures and their predicted function simplified the impacts from natural processes and anthropogenic activities in coastal areas. Mar Pollut Bull 131:481–495. 10.1016/j.marpolbul.2018.04.052

83. Tamim U, Khan R, Jolly YN, Fatema K, Das S, Naher K, Islam MA, Islam SMA, Hossain SM. (2016). Elemental distribution of metals in urban river sediments near an industrial effluent source. Chemosphere 155, 509–518. 10.1016/j.chemosphere.2016.04.09 9.

84. Tiquia, S.M., 2010. Metabolic diversity of the heterotrophic microorganisms and potential link to pollution of the Rouge River. J. Environ. Pollut. 158, 1435–1443. 10.1016/j.envpol.2009.12.035

85. Wang, L.; Zheng, B.; Nan, B.; Hu, P. (2014). Diversity of bacterial community and detection of nirS—And nirK-encoding denitrifying bacteria in sandy intertidal sediments along Laizhou Bay of Bohai Sea, China. Mar. Pollut. Bull. 88, 215–223. 10.1016/j.marpolbul.2014.09.002

86. Wang, L., Zhang, J., Li, H., Yang, H., Peng, C., Peng, Z., & Lu, L. (2018). Shift in the microbial community composition of surface water and sediment along an urban river. Science of the Total Environment, 627, 600–612. 10.1016/j.scitotenv.2018.01.203

87. Wang, W.; Yi, Y.; Yang, Y.; Zhang, S.; Zhou, Y.; Wang, X.; Yang, Z. (2020). Impact of anthropogenic activities on the sediment microbial communities of Baiyangdian shallow lake. Int. J. Sediment Res., 35, 180–192. DOI: 10.1016/j.ijsrc.2019.10.006

88. Wang, F., Dong, W., Zhao, Z., Wang, H., Li, W., Chen, G., Wang, F., Zhao, Y., Huang, J., Zhou, T. (2021). Heavy metal pollution in urban river sediment of different urban functional areas and its influence on microbial community structure. Sci. Total Environ. 778, 146383 10.1016/j.scitotenv.2021.146383.

89. Wemheuer, F., Taylor, J. A., Daniel, R., Johnston, E., Meinicke, P., Thomas, T., et al. (2020). Tax4Fun2: prediction of habitat-specific functional profiles and functional redundancy based on 16S rRNA gene sequences. Environ. Microbiome 15:11. 10.1186/s40793-020-00358-7

90. Wen, Y., Xiao, M., Chen, Z., Zhang, W., & Yue, F. (2023). Seasonal Variations of Dissolved Organic Matter in Urban Rivers of Northern China. Land, 12(2), 1–18. 10.3390/land12020273

91. Wéry, N., Lhoutellier, C., Ducray, F., Delgenès, J.-P., Godon, J.-J. (2008). Behaviour of pathogenic and indicator bacteria during urban wastewater treatment and sludge composting, as revealed by quantitative PCR. J. Water Res. 42, 53–62. 10.1016/j.watres.2007.06.048

92. Withers, P., Jarvie, H., (2008). Delivery and cycling of phosphorus in rivers: a review. Sci. Total Environ. 400, 379–395. 10.1016/j.scitotenv.2008.08.002

93. Wu, C., Narale, D. D., Cui, Z., Wang, X., Liu, H., Xu, W., Zhang, G., & Sun, J. (2022). Diversity, structure, and distribution of bacterioplankton and diazotroph communities in the Bay of Bengal during the winter monsoon. Frontiers in Microbiology, 13. 10.3389/fmicb.2022.987462

94. Wu, X., Wu, L., Liu, Y., Zhang, P., Li, Q., Zhou, J., et al., (2018). Microbial Interactions With Dissolved Organic Matter Drive Carbon Dynamics and Community Succession. Front. Microbiol. 9:1234. doi: 10.3389/fmicb.2018.01234

95. Wu, H., Li, Y., Zhang, W., Wang, C., Wang, P., Niu, L., Du, J., Gao, Y., (2019). Bacterial community composition and function shift with the aggravation of water quality in a heavily polluted river. J. Environ. Manage. 237, 433–441. 10.1016/j.jenvman.2019.02.101.

96. Xie, G., Tang, X., Shao, K., Zhu, G., & Gao, G. (2021). Bacterial diversity, community composition and metabolic function in Lake Tianmuhu and its dammed river: Effects of domestic wastewater and damming. Ecotoxicology and Environmental Safety, 213, 112069. 10.1016/j.ecoenv.2021.112069

97. Xiong, W., Sun, Y., Zhang, T., Ding, X., Li, Y., Wang, M., et al., (2015). Antibiotics, antibiotic resistance genes, and bacterial community composition in fresh water aquaculture environment in China. Microb. Ecol. 70, 425e432. 10.1007/s00248-015-0583-x

98. Yin, H., Niu, J., Ren, Y., Cong, J., Zhang, X., Fan, F., et al. (2015). An integrated insight into the response of sedimentary microbial communities to heavy metal contamination. Sci. Rep. 5. 10.1038/srep14266

99. Zhang J, Zhang X, Liu Y, Xie S, Liu Y. (2014). Bacterioplankton communities in a high-altitude freshwater wetland. Ann Microbiol. 64(3):1405–1411. 10.1007/s13213-013-0785-8

100. Zhang, L., Li, L., Liu, M., Hu, Y., Jiang, J. (2019a). Temporal and spatial variations of bacterial community compositions in two estuaries of Chaohu Lake. J. Oceanol. Limnol. 1e14. 10.1007/s00343-019-9096-7

101. Zhang, M., Wu, Z., Sun, Q., Ding, Y., Ding, Z., Sun, L. (2019b). Response of chemical properties, microbial community structure and functional genes abundance to seasonal variations and human disturbance in Nanfei River sediments. Ecotoxicol. Environ. Saf. 183. 10.1016/j.ecoenv.2019.109601

102. Zhang, L., Zhao, F., Li, X., Lu, W. (2020a). Contribution of influent rivers affected by different types of pollution to the changes of benthic microbial community structure in a large lake. Ecotoxicol. Environ. Saf. 198, 110657 10.1016/j.ecoenv.2020.110657.

103. Zhang, W., Wang, H., Li, Y., Lin, L., Hui, C., Gao, Y., Niu, L., Zhang, H., Wang, L., Wang, P., Wang, C. (2020b). Bend-induced sediment redistribution regulates deterministic processes and stimulates microbial nitrogen removal in coarse sediment regions of the river. Water Res. 170, 115315 10.1016/j.watres.2019.115315.

104. Zhang, L., Zhou, Y., Cheng, Y., Lu, W., & Liang, Y. (2021). Effect of different types of industrial wastewater on the bacterial community of urban rivers. Journal of Freshwater Ecology, 36(1), 31–48. 10.1080/02705060.2021.1871978

105. Zhao, Y., Xia, X. H., Yang, Z. F., & Wang, F. (2012). Assessment of water quality in Baiyangdian Lake using multivariate statistical techniques. Procedia Environmental Sciences, 13, 1213–1226. 10.1016/j.proenv.2012.01.115

106. Zheng, G., Yampara-Iquise, H., Jones, J., Andrew Carson, C., 2009. Development of Faecalibacterium 16S rRNA gene marker for identification of human faeces. J. Appl. Microbiol. 106, 634–641. 10.1111/j.1365-2672.2008.04037.x

107. Zhu, C., Zhang, J., Nawaz, M.Z., Mahboob, S., Al-Ghanim, K.A., Khan, I.A., et al. (2019). Seasonal succession and spatial distribution of bacterial community structure in a eutrophic freshwater Lake, Lake Taihu. Sci. Total Environ. 669, 29e40. 10.1016/j.scitotenv.2019.03.087

