## Supplemental file for "Temporal and spatial distribution of microbial community in urban river in Dhaka"

### Supplementary Data

**Table S1**

Details of sampling stations.

| Station Name | Station code | Coordinate | |
|---|---|---|---|
| | | **Latitude** | **Longitude** |
| Mirpur Boro Bazar Jame Mosque (Turag River) | 01TR | 23.789277 | 90.338991 |
| Begunbari Down Field (Karnatali River) | 02KR | 23.791279 | 90.331836 |
| Amin Bazar Bridge (Junction of Three River) | 03BR | 23.784321 | 90.336029 |
| Gabtoli BalurGhat Mosque | 04BR | 23.776119 | 90.333733 |
| Dhaka Uddan Housing River View Point | 05BR | 23.757947 | 90.335353 |
| Bosila Bridge | 06BR | 23.745428 | 90.344710 |
| Gudaraghat Jhauchor Jame Mosque | 07BR | 23.727716 | 90.357684 |
| Nababchor Bridge (Canal side) | 08CS | 23.721950 | 90.353932 |
| Mandail Mosque | 09BR | 23.708223 | 90.382817 |
| Lohar Bridge (Canal side) | 10CS | 23.712486 | 90.385267 |
| Mitford Ghat | 11BR | 23.710263 | 90.400170 |
| Sadar Ghat | 12BR | 23.702048 | 90.414199 |
| Postagola Cantonment (B-C Friendship Bridge) | 13BR | 23.689226 | 90.426215 |

***KR = Karnatali River; BR = Buriganga River; TR = Turag River and CS = Canal Side***

**Table S2**

List of physicochemical water quality parameters and analytical methods.

| S.l | Parameters Name | Unit | Analysis Method |
|---|---|---|---|
| 01 | pH | … | pH meter (HACH_HQ11d) |
| 02 | Turbidity | NTU | Turbidity meter (HACH_HQ11d) |
| 03 | Electric Conductivity (EC) | $\mu$S/cm | Conductivity meter (Multiline_p4) |
| 04 | Total Dissolved Solid (TDS) | mg/l | Gravimetric method |
| 05 | Temperature | $^{o}$C | pH meter (HACH_HQ11d) |
| 06 | Dissolved Oxygen (DO) | mg/l | DO meter (HACH_HQ11d) |
| 07 | Biochemical Oxygen Demand ($BOD_5$) | mg/l | 5 Days Incubation |
| 08 | Chemical Oxygen Demand (COD) | mg/l | Spectrophotometric (HACH_DR 6000) |
| 09 | Nitrogen Di-oxide ($NO_2$) | mg/l | Spectrophotometric (HACH_DR 6000) |
| 10 | Ammonium Nitrate ($NH_3$-N) | mg/l | Spectrophotometric (HACH_DR 6000) |
| 11 | Total Phosphorus (TP) | mg/l | Spectrophotometric (HACH_DR 6000) |
| 12 | Chloride ($Cl^-$) | mg/l | Spectrophotometric (HACH_DR 6000) |
| 13 | Lead (pb) | ppm | Atomic Absorption Spectrophotometer (SHIMADZU_AA-7000) |
| 14 | Cadmium (Cd) | ppm | Atomic Absorption Spectrophotometer (SHIMADZU_AA-7000) |
| 15 | Chromium (Cr) | ppm | Atomic Absorption Spectrophotometer (SHIMADZU_AA-7000) |
| 16 | Cupper (Cu) | ppm | Atomic Absorption Spectrophotometer (SHIMADZU_AA-7000) |
| 17 | Nickel (Ni) | ppm | Atomic Absorption Spectrophotometer (SHIMADZU_AA-7000) |
| 18 | Zinc (Zn) | ppm | Atomic Absorption Spectrophotometer (SHIMADZU_AA-7000) |

**Table S3**

Sequence Count in every sample

| Sample ID | Forward Sequence Count | Reverse Sequence Count |
|---|---|---|
| BR42 | 126954 | 126954 |
| TR40 | 121911 | 121911 |
| BR46 | 118881 | 118881 |
| KR41 | 117481 | 117481 |
| BR44 | 117470 | 117470 |
| BR3 | 117388 | 117388 |
| CS49 | 116894 | 116894 |
| BR35 | 116860 | 116860 |
| BR52 | 116086 | 116086 |
| BR50 | 112555 | 112555 |
| TR1 | 110894 | 110894 |
| BR51 | 108340 | 108340 |
| BR25 | 107941 | 107941 |
| BR48 | 107274 | 107274 |
| BR24 | 106956 | 106956 |
| CS8 | 106793 | 106793 |
| BR4 | 106660 | 106660 |
| BR6 | 104786 | 104786 |
| BR5 | 104270 | 104270 |
| CS21 | 102984 | 102984 |
| KR28 | 102723 | 102723 |
| BR7 | 102034 | 102034 |
| BR22 | 101612 | 101612 |
| BR38 | 101398 | 101398 |
| TR14 | 100137 | 100137 |
| KR2 | 100114 | 100114 |

| | | |
|---|---|---|
| BR18 | 99750 | 99750 |
| BR9 | 99747 | 99747 |
| BR19 | 99666 | 99666 |
| BR33 | 99609 | 99609 |
| BR31 | 99489 | 99489 |
| BR11 | 98700 | 98700 |
| BR13 | 98534 | 98534 |
| CS23 | 98379 | 98379 |
| BR20 | 98243 | 98243 |
| BR16 | 98204 | 98204 |
| BR26 | 98113 | 98113 |
| BR12 | 98092 | 98092 |
| CS10 | 97901 | 97901 |
| BR37 | 97135 | 97135 |
| BR32 | 95911 | 95911 |
| BR17 | 91755 | 91755 |
| BR30 | 90711 | 90711 |
| BR45 | 72575 | 72575 |
| BR43 | 67229 | 67229 |
| KR15 | 64691 | 64691 |
| BR39 | 64631 | 64631 |
| CS36 | 62615 | 62615 |
| CS34 | 50674 | 50674 |
| BR29 | 41521 | 41521 |
| TR27 | 15480 | 15480 |
| | **Total** | **4956751** |
| | **Average (Per sample)** | **97,191** |

**Table S4**

List of **Phyla** among the four different seasons with proportion of relative abundance.

| Phylum | Pre-monsoon | Monsoon | Post-monsoon | Winter | Average |
|---|---|---|---|---|---|
| Actinobacteria | 13.5 | 18.39 | 6.26 | 7.95 | **11.53** |
| Bacteroidetes | 3.65 | 4.38 | 10.05 | 7.2 | **6.32** |
| Cyanobacteria | 4.35 | 9.58 | 2.82 | 2.69 | **4.86** |
| Firmicutes | 12.09 | 7.25 | 12.39 | 13.6 | **11.33** |
| Proteobacteria | 66.41 | 60.4 | 67.49 | 68.57 | **65.72** |

**Table S5**

List of **Genera** among the four different seasons with the proportion of relative abundance.

| Genus_Proportion of Relative Abundance greater than 3% | | | | | |
|---|---|---|---|---|---|
| **Genera** | **Pre-monsoon** | **Monsoon** | **Post-monsoon** | **Winter** | **Average** |
| Acinetobacter | 21.32 | 4.97 | 34.03 | 22.98 | 20.82 |
| Arcobacter | 7.65 | 0.00 | 27.58 | 11.66 | 11.72 |
| C39 | 11.61 | 16.34 | 12.78 | 9.11 | 12.46 |
| Dechloromonas | 9.11 | 12.59 | 6.55 | 4.10 | 8.09 |
| Hydrogenophaga | 7.18 | 5.42 | 7.48 | 6.87 | 6.74 |
| Klebsiella | 6.43 | 0.00 | 0.00 | 0.00 | 1.61 |
| Limnohabitans | 4.31 | 6.96 | 0.00 | 0.00 | 2.82 |
| Novosphingobium | 15.98 | 16.42 | 0.00 | 5.27 | 9.42 |
| Planktothrix | 5.49 | 0.00 | 0.00 | 0.00 | 1.37 |
| Hylemonella | 0.00 | 3.30 | 0.00 | 0.00 | 0.82 |
| Methylocaldum | 0.00 | 3.75 | 0.00 | 0.00 | 0.94 |
| Methylosinus | 0.00 | 4.70 | 0.00 | 0.00 | 1.18 |
| Synechococcus | 0.00 | 17.42 | 0.00 | 0.00 | 4.36 |
| Polynucleobacter | 0.00 | 3.91 | 0.00 | 0.00 | 0.98 |
| Oscillospira | 0.00 | 0.00 | 4.56 | 0.00 | 1.14 |
| Ralstonia | 0.00 | 0.00 | 7.01 | 0.00 | 1.75 |
| Lactobacillus | 0.00 | 0.00 | 0.00 | 3.80 | 0.95 |
| Rothia | 0.00 | 0.00 | 0.00 | 3.10 | 0.77 |

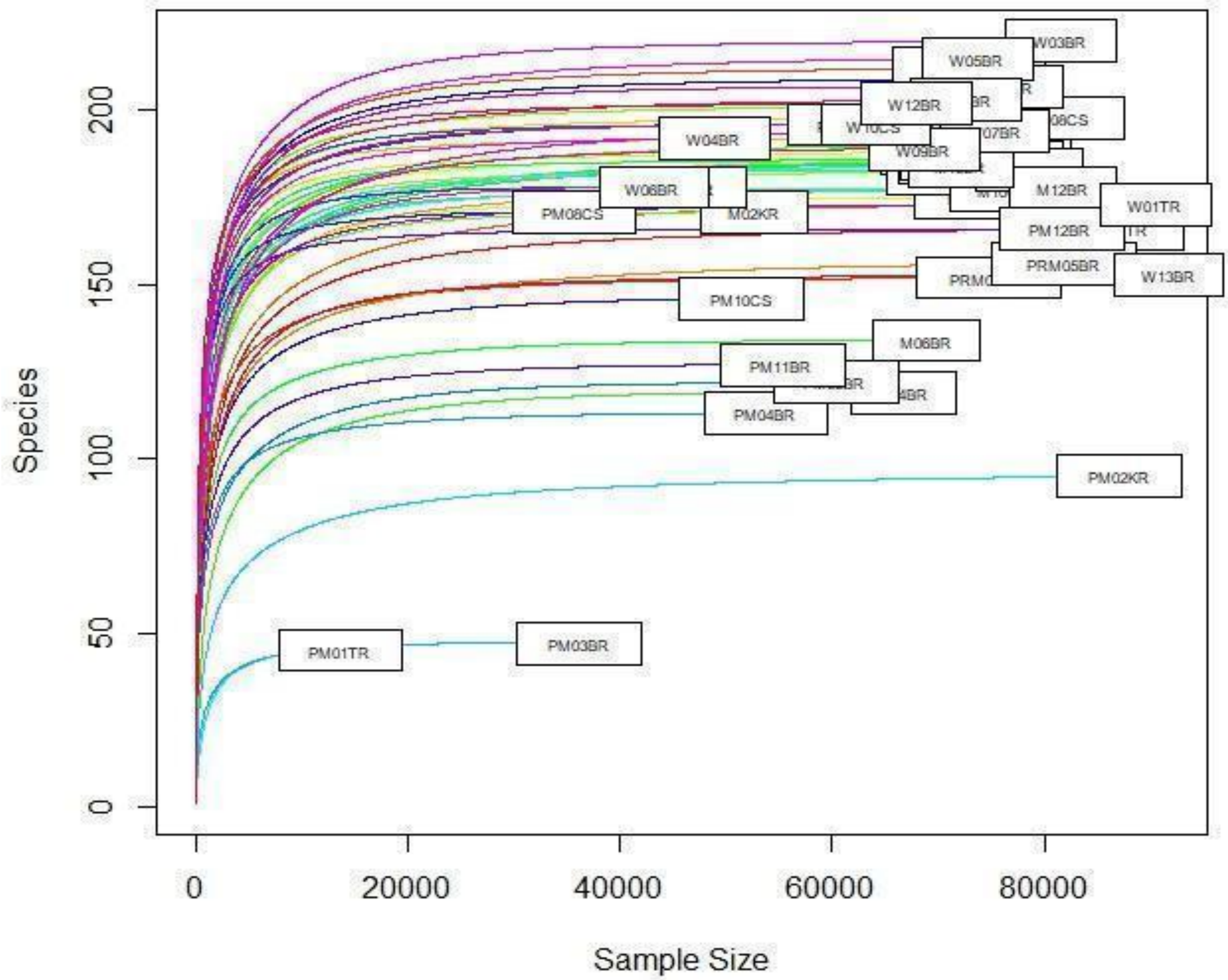


**Figure S1.** Rarefaction curve which illustrates whether a sample has been sequenced to an extent sufficient to represent its true diversity or not.

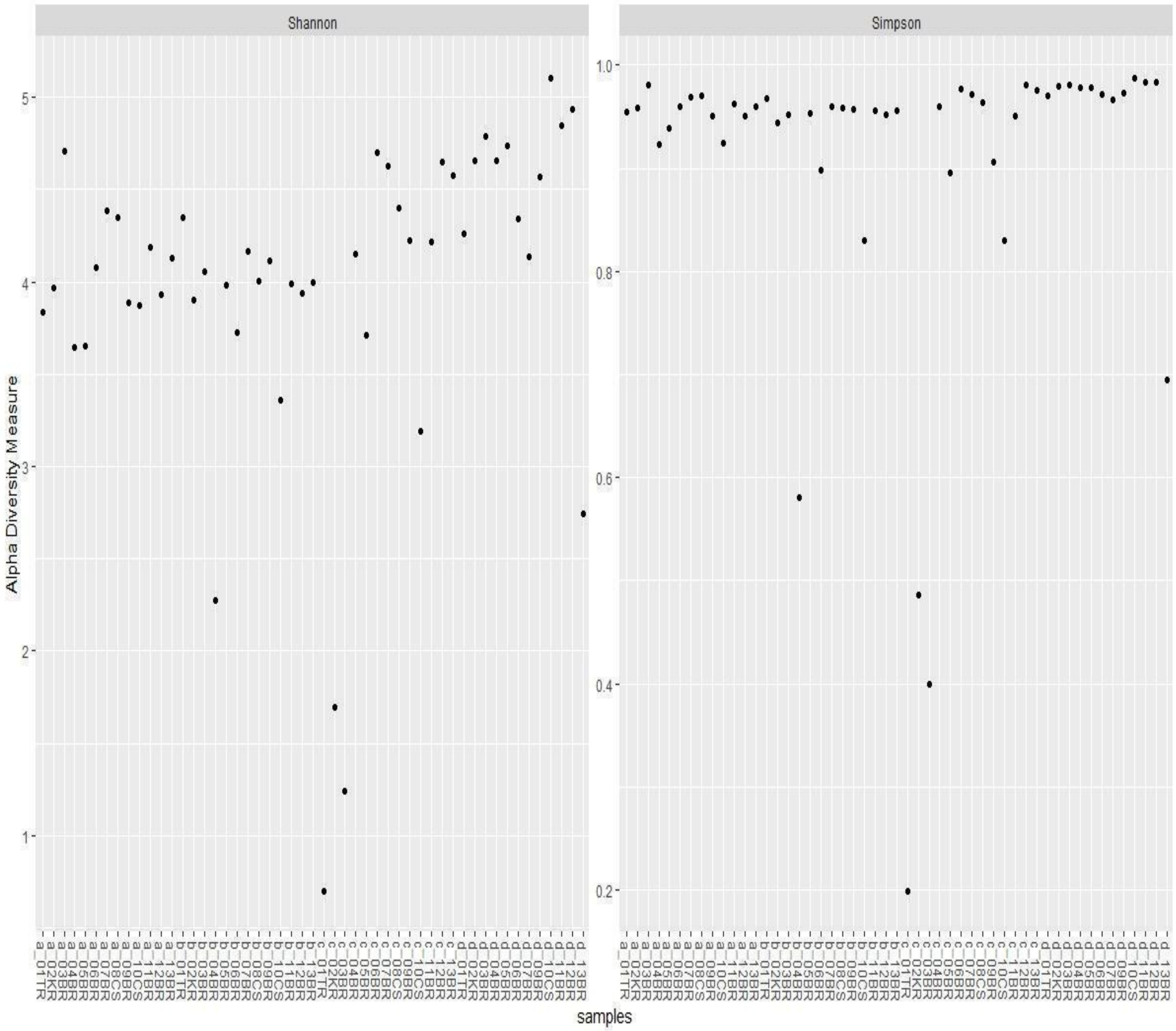


**Figure S2.** Alpha diversity (Shannon & Simpson) at individual sampling points. where a, b, c and d indicate Pre-monsoon, Monsoon, Post-monsoon and Winter respectively.
